# Glycolytic reprogramming fuels myeloid cell-driven hypercoagulability

**DOI:** 10.1101/2023.04.20.537683

**Authors:** Aisling M. Rehill, Gemma Leon, Sean McCluskey, Ingmar Schoen, Yasmina Hernandez-Santana, Stephanie Annett, Paula Klavina, Tracy Robson, Annie M. Curtis, Thomas Renné, Seamus Hussey, James S. O’Donnell, Patrick T. Walsh, Roger J.S. Preston

**Affiliations:** Irish Centre for Vascular Biology, School of Pharmacy and Biomolecular Sciences, RCSI University of Medicine and Health Sciences, Dublin, Ireland; National Children’s Research Centre, Children’s Health Ireland Crumlin, Dublin, Ireland; Dept of Clinical Medicine, Trinity Translational Medicine Institute, Trinity College Dublin, Ireland; Institute of Clinical Chemistry and Laboratory Medicine, University Medical Center Hamburg-Eppendorf, Hamburg, Germany; Department of Paediatrics, University College Dublin and RCSI University of Medicine and Health Sciences, Dublin, Ireland

**Keywords:** coagulation, fibrinolysis, inflammation, macrophages, protein C

## Abstract

**Background:** Myeloid cell metabolic reprogramming is a hallmark of inflammatory disease, however, its role in inflammation-induced hypercoagulability is poorly understood.

**Objective/Methods:** Using novel myeloid cell-based global haemostasis assays and murine models of immunometabolic disease, we evaluated the role of inflammation-associated metabolic reprogramming in regulating blood coagulation.

**Results:** Glycolysis was essential for enhanced activated myeloid cell tissue factor expression and decryption, driving increased cell-dependent thrombin generation in response to inflammatory challenge. Similarly, inhibition of glycolysis enhanced activated macrophage fibrinolytic activity via reduced plasminogen activator inhibitor 1 (PAI-1)-activity. Macrophage polarisation or activation markedly increased endothelial protein C receptor (EPCR) expression on monocytes and macrophages, leading to increased myeloid cell-dependent protein C activation. Importantly, inflammation-dependent EPCR expression on tissue-resident macrophages was also observed *in vivo*. Adipose tissue macrophages from obese mice fed a high-fat diet exhibited significantly enhanced EPCR expression and APC generation compared to macrophages isolated from the adipose tissue of healthy mice. Similarly, the induction of colitis in mice prompted infiltration of EPCR^+^ innate myeloid cells within inflamed colonic tissue that were absent from the intestinal tissue of healthy mice.

**Conclusion:** Collectively, this study identifies immunometabolic regulation of myeloid cell hypercoagulability, opening new therapeutic possibilities for targeted mitigation of thrombo-inflammatory disease.

**ESSENTIALS:**

- Inflammation-mediated glycolytic reprogramming enables myeloid cell-induced hypercoagulability and antifibrinolytic activity.
- 2-Deoxy-D-glucose (2-DG) inhibits the expression of transcription factors necessary for inflammation-induced procoagulant gene expression.
- Myeloid cell membrane regulation of tissue factor procoagulant activity is glycolysis-dependent.
- Activation of myeloid innate immunity dysregulates activated protein C anticoagulant pathway activity.

## INTRODUCTION

Monocytes and macrophages play an important role in thrombus formation during inflammatory disease. Enhanced myeloid cell tissue factor (TF) expression occurs in response to pro-inflammatory stimuli and has been reported in many inflammatory disease states associated with an elevated risk of venous thromboembolism (VTE), including obesity [1,2], sickle cell disease [3,4], inflammatory bowel disease [5,6] and chronic myeloproliferative disorders [7,8]. Myeloid cells can also regulate the rate of clot fibrinolysis through secretion of plasminogen activator inhibitor 1 (PAI-1) in response to infection or inflammatory challenges. Myeloid cell-derived PAI-1 is proposed to directly contribute to many immuno-thrombotic conditions [9] and high PAI-1 levels have been reported in individuals with obesity [10], sepsis [11,12] and severe COVID-19 [13,14]. In contrast to the established role of immune cell-mediated coagulation activation in response to inflammatory stimuli, its impact on endogenous anticoagulant activity is poorly understood. The protein C pathway serves a critical role in regulating thrombin generation and fibrin deposition. Zymogen plasma protein C conversion to activated protein C (APC) by the thrombin-thrombomodulin (TM) complex is accelerated 4-20-fold by protein C binding to EPCR [15,16] and is therefore regulated by EPCR and TM membrane expression. Once APC has been generated, it must detach from EPCR to localise on anionic phospholipid surfaces to degrade activated coagulation cofactors factor V (FVa) and factor VIII (FVIIIa) and limit further thrombin generation. APC also initiates anti-inflammatory myeloid cell signalling via distinct, cell-specific receptor pathways that exert significant anti-inflammatory benefits in both acute and chronic inflammatory disease models [17–19]. EPCR and TM are abundantly expressed on endothelial cells but also on human and mouse myeloid cell populations [20–23]. However, little is known about the impact of macrophage activation or phenotype in modulating APC activities.

Upon activation, myeloid cells undergo extensive metabolic re-programming to enable increased cytokine biosynthesis and phagocytic activity [24]. Specifically, activated (or ‘M1’) macrophages exhibit increased glycolysis and pentose phosphate pathway activity [25]. In contrast, anti-inflammatory (or ‘M2’) macrophages exhibit distinct metabolic features that promote the resolution of inflammation [26]. *In vivo*, tissue-resident macrophages also exhibit altered metabolism during inflammation. Previous studies have highlighted a critical role for glycolytic enzymes in thrombo-inflammatory disorders, such as stroke and atherosclerosis [27–29]. For example, pyruvate kinase muscle 2 (PKM2) promotes neutrophil activation and thrombo-inflammatory cytokine generation [30] and myeloid-specific PKM2 inhibition is associated with reduced infarct size and improved outcomes in ischemic stroke [27,30,31]. Despite this, the role of cell metabolism in regulating myeloid cell-dependent hypercoagulability is poorly understood.

In this study, we demonstrate that glycolytic reprogramming of monocyte/macrophages in response to pro-inflammatory stimulation directly influences myeloid cell-driven hypercoagulability. In particular, we show that enhanced glycolysis is necessary for increased myeloid cell TF expression and decryption and enhanced production of PAI-1/2 to limit fibrinolysis. Furthermore, we demonstrate that myeloid cell metabolic reprogramming also alters EPCR-dependent APC generation. These data suggest that targeted inhibition of myeloid cell metabolism may represent a novel approach to limit procoagulant activity during inflammation.

## MATERIALS AND METHODS

Further details can be found in the *‘Supplemental Methods and Materials’*.

### Generation of bone marrow-derived macrophages (BMDMs)

Bone marrow-derived macrophages (BMDMs) were isolated and polarised, as previously described [32,33]. For detailed protocol, see ‘*Supplementary Materials’*. BMDMs were polarised towards M1 using 20ng/mL IFNγ and 100ng/mL LPS and towards M2 using 20ng/mL IL-4 and IL-13. BMDMs were polarised for 6 hours before RNA isolation and for 18 hours before functional assays and ELISAs. Cells were treated with 5µg/mL ssRNA for 12 hours before RNA isolation and 24-48 hours before functional assays.

### Isolation and culture of human monocytes

Anonymised healthy donor buffy coats were obtained from the Irish Blood Transfusion Service, St. James’ Hospital, Dublin and monocytes were isolated as previously described [34]. Monocytes were activated with 100ng/mL LPS for 18 hours before functional assays. Cells were treated with 5µg/mL ssRNA for 24-48 hours before functional assays.

### Myeloid cell-based thrombin generation assay

Unless otherwise stated, all reagents were obtained from Stago. Cells were seeded onto a 96-well plate (5 × 10^4^ cells per well for macrophages and 2 × 10^5^ cells per well for human monocytes) and left overnight to adhere. Following cell stimulations, supernatants were removed, and cells were washed once with PBS. 20µL MP-reagent was added to 80µL factor XII (FXII)-deficient plasma (HTI). To examine the procoagulant activity of each cell releasate, 80µL of supernatants were added to plasma and MP reagent. The assay was initiated using 20µL FluCa-kit. Fluorescence was quantified using Thrombinoscope software on a Fluoroskan fluorometer.

### Myeloid cell-based plasmin generation assay

Macrophages were seeded onto 96-well plates (5 × 10^4^ cells per well). Following treatment, macrophages were washed with PBS and then incubated with 100µL serum-free DMEM supplemented with 1U/mL penicillin, 0.1mg/mL streptomycin, 50ng/mL t-PA and 400nM Glu-plasminogen (HTI) for 1.5 hours. Supernatants were then transferred to a black flat-bottomed 96-well plate. 20µL Boc-Glu-Lys-Lys fluorometric substrate was added to each well to enable plasmin detection (final concentration, 1260µM; Bachem). Substrate was diluted in 34mM CaCl_2_ containing TBS. Assays were performed with each treatment group in duplicate per assay and at least biological triplicates, with a negative control well containing only serum-free DMEM. Fluorescence was quantified using the Ascent software for Fluoroskan Ascent Fluorometer. Plasmin generation at 60 minutes (PG^60^) was assessed for each assay condition.

### mRNA isolation and RT-qPCR

5 × 10^5^ cells/mL were lysed using 350µL lysis buffer and total RNA isolated according to manufacturer’s instructions (Ambion PureLink RNA isolation kit). Gene expression was determined by performing RT-qPCR in duplicate with PowerUp^TM^ SYBR^TM^ green master mix (Thermofisher) using a 7500 Fast system (Applied Biosystems). Relative-fold changes in mRNA expression were calculated using the cycle threshold (C_T_) and normalised to the *RPS18* housekeeping gene. Primer sequences are listed in **Supplementary Table 1**.

### Macrophage siRNA transfection

SMARTpool siRNA guides were designed by Horizon Discovery and chosen using the siGENOME siRNA search tool. siRNAs (20nM) were transfected using Lipofectamine RNAiMax (Invitrogen). For detailed protocol, see ‘*Supplementary Materials’*.

### Protein C activation assay

Cells were seeded onto a 96-well plate (1 × 10^5^ cells per well for macrophages and 4 × 10^5^ cells per well for human monocytes) and left overnight to adhere. Cells were incubated with 100nM human or mouse PC (HTI and Sino Biologicals, respectively) and 5nM thrombin (HTI) in phenol-free DMEM with 3mM CaCl_2_ and 0.6mM MgCl_2_ for 1 hour at 37°C. The reaction was stopped with a 10-fold molar excess of hirudin (Sigma) and APC was detected using an APC chromogenic substrate (Quadratech).

### Confocal microscopy

Murine bone marrow-derived macrophages grown on gelatin-coated coverslips were fixed and permeabilised with 100% ice-cold methanol for 10 minutes at −20°C. Cells were washed with PBS and stored at 4°C until ready for staining. Cells were probed with a primary anti-mouse ASMase antibody (2μg; ThermoFisher Scientific) overnight at 4°C and then labelled with Invitrogen Goat anti-Rabbit IgG (H+L) Highly Cross-Adsorbed Secondary Antibody, Alexa Fluor Plus 488 (0.4ug; ThermoFisher Scientific) for 1 hour at room temperature. Cells were then washed, stained with CellMask™ Deep Red Plasma Membrane Stain (1000X; ThermoFisher Scientific) for 10 minutes at 37°C, and mounted onto glass slides containing one drop of SlowFade™ Gold Antifade Mountant with DAPI (Thermofisher Scientific). Slides were imaged using a Zeiss LSM700 confocal microscope and analysed using FIJI open-source image analysis software platform[35]. Data is representative of 4 independent experiments. Corrected total cell fluorescence (CTCF) was measured in 92-99 cells per condition (M0 = 92, M1 = 98, M1+2DG = 99).

### Flow cytometry

All fluorescently labelled antibodies used were purchased from eBiosciences™ (anti-F480, anti-EPCR, anti-TM, anti-CD45, anti-MHC-II, anti-LY6C, anti-CD11b, anti-CD11c, anti-CD14, anti-CD86 and anti-CD206) except for anti-TF antibody (AF3178; R&D Systems). Cell viability was measured using the LIVE/DEAD™ Fixable Aqua Dead Cell Stain Kit (ThermoFisher Scientific), and cell numbers were determined using CountBright™ Absolute Counting Beads (ThermoFisher Scientific). Before staining, cells were first incubated with an anti-CD16/CD32 Monoclonal Antibody to block FC receptors (Invitrogen). Cells were washed using PBS and incubated with LIVE/DEAD™ dye for 30 minutes at 4°C. Cells were washed using PBS supplemented with 2% FBS and incubated with fluorescently labelled antibodies for 1 hour at 4°C. Fluorescence Minus One (FMO) controls were used to assess positive staining. Multi-parameter analysis was then performed on an LSR/Fortessa (BD) and analysed using FlowJo 10 software.

### Protein binding assays

Recombinant mouse lactadherin (R&D Systems) and human APC (Cambridge ProteinWorks) were fluorescently labelled using Lightning-Link® Rapid Alexa Fluor 488 Antibody Labelling Kits in accordance with the manufacturer’s instructions (Novus Biologicals). Macrophages were incubated with the labelled proteins or anti-mouse F4/80 (ThermoFisher Scientific) for 2 hours. Multi-parameter flow cytometry was performed on an LSR/Fortessa (BD) and analysed using FlowJo™ 10 software to detect cell surface protein binding.

### Dextran sulfate sodium (DSS) induction of colitis in mice

Colitis was induced in wild-type C57/BL6 mice aged 8-12 weeks by adding 2% (w/v) DSS to drinking water. Mice were monitored daily for five days, and their weight loss, stool consistency and rectal bleeding were recorded and used to calculate Disease Activity Index. Previously co-housed age-matched wild-type C57/BL6 mice were kept on normal drinking water to act as a control. Animals were housed under specific pathogen-free (SPF) conditions in a temperature-controlled unit with a 12-h light/dark cycle at the Comparative Medicine Unit (CMU) in Trinity Translational Medicine Institute (TTMI), St. James Hospital (Dublin). Water and food were provided *ad libitum*. All animal experiments were approved by the Health Products Regulatory Authority under Project License AE19136/P125. Mouse lamina propria leukocyte populations were isolated as previously described [36], and isolated leukocytes were analysed by flow cytometry (full protocol in *‘Supplementary Methods’*).

### High-fat diet (HFD) induced obesity model

Male C57BL/6nCrl mice aged 7-8 weeks old were fed a 60% high-fat diet (HFD, F3282, Datesend) or standard chow (Verified 75 IF irradiated, Lab Supply) for 12 weeks. Mouse weights were measured weekly. C57BL/6nCrl mice were obtained from Charles Rivers (UK) and were housed in the Biological Resource Unit at RCSI University of Medicine and Health Sciences, Dublin, Ireland. All animal experiments were approved by the Health Products Regulatory Authority under Project License AE19127/P045 and Individual License AE19127/P045. When HFD mice reached ∼45g, animals were sacrificed as described above, and the white adipose tissue (WAT) from inguinal tissue was collected, as previously described [37] (detailed protocol in ‘*Supplementary Methods’*). Macrophages were isolated by magnetic cell isolation technology using CD11b microbeads (Miltenyi Biotec), according to the manufacturer’s protocol. Flow cytometry confirmed ATMs as a macrophage population by analysing F4/80 and CD11b expression.

### Statistical analysis

Data were analysed for normal distribution (Shapiro–Wilk or Kolmogorov–Smirnov test) and equality of variance (F test). One-way ANOVA analysis was used to compare the means of 3 or more groups. Where significance was found in the ANOVA analysis, Tukey’s multiple comparison test was used to define differences between individual groups. To compare the mean of the two groups, a Student’s t-test was used, unpaired for *in vivo* experiment analysis and paired for *in vitro* experiment analysis or Mann Whitney U test for nonparametric data, as appropriate. All analysis and graph representation were performed using GraphPad Prism 9 software, and data are shown as mean ± SD.

## RESULTS

### Glycolysis is essential for pro-inflammatory myeloid cell procoagulant activity

Murine bone marrow-derived macrophages (BMDMs) were polarised towards M1 or M2 phenotypes **(Supplementary Figure 1a-d)**. Initial characterisation of M1- and M2-polarised BMDMs revealed distinct expression profiles for genes encoding haemostasis-related proteins **(Supplementary Figure 1e)**. In M1 BMDMs, *PROCR* (257-fold), *F10* (34-fold) and *F3* (8-fold) expression were significantly up-regulated. In contrast, except for slightly increased *F10* and *F7* expression, haemostasis-associated genes were not significantly impacted by M2 polarisation. We developed a novel myeloid cell-based calibrated automated thrombinography (CAT) assay to investigate immune cell-dependent hypercoagulability in plasma. We used FXII-deficient plasma (to prevent contact pathway activation) incubated with BMDMs **(Figure 1a)**. M1-polarised BMDMs promoted significantly faster thrombin generation compared to either naïve or M2 BMDMs, such that M1 BMDMs reduced lag-time by 7 minutes (∼50%) compared to naïve BMDMs **(Figure 1b-c)**. Thrombin generation was TF-dependent, as minimal thrombin generation was observed in FVII-deficient plasma **(Supplementary Figure 1f-i)**. Increased *F3* gene expression was also observed in both M1 polarised THP1-derived human macrophages and primary human monocyte-derived macrophages (HMDMs) **(Supplementary Figure 2a-d)**. To confirm that this modified CAT assay could also detect hypercoagulability in human myeloid cells, the assay was performed with polarised HMDMs differentiated with both human serum **(Supplementary Figure 2e-h)** or in the presence of human M-CSF **(Supplementary Figure 2i-l)**. A similar decrease in lagtime was observed in the presence of M1 polarised HMDMs, while M2 HMDMs exhibited similar procoagulant activity as naive macrophages **(Supplementary Figure 2f and j)**. There was no observed difference when HMDMs were differentiated using human serum or M-CSF. Additionally, human peripheral blood monocytes were isolated from healthy donors and activated using LPS. LPS treatment of human monocytes decreased lag-time by 8 minutes compared to naïve monocytes **(Figures 1d and 1e)**.

**Figure 1:**
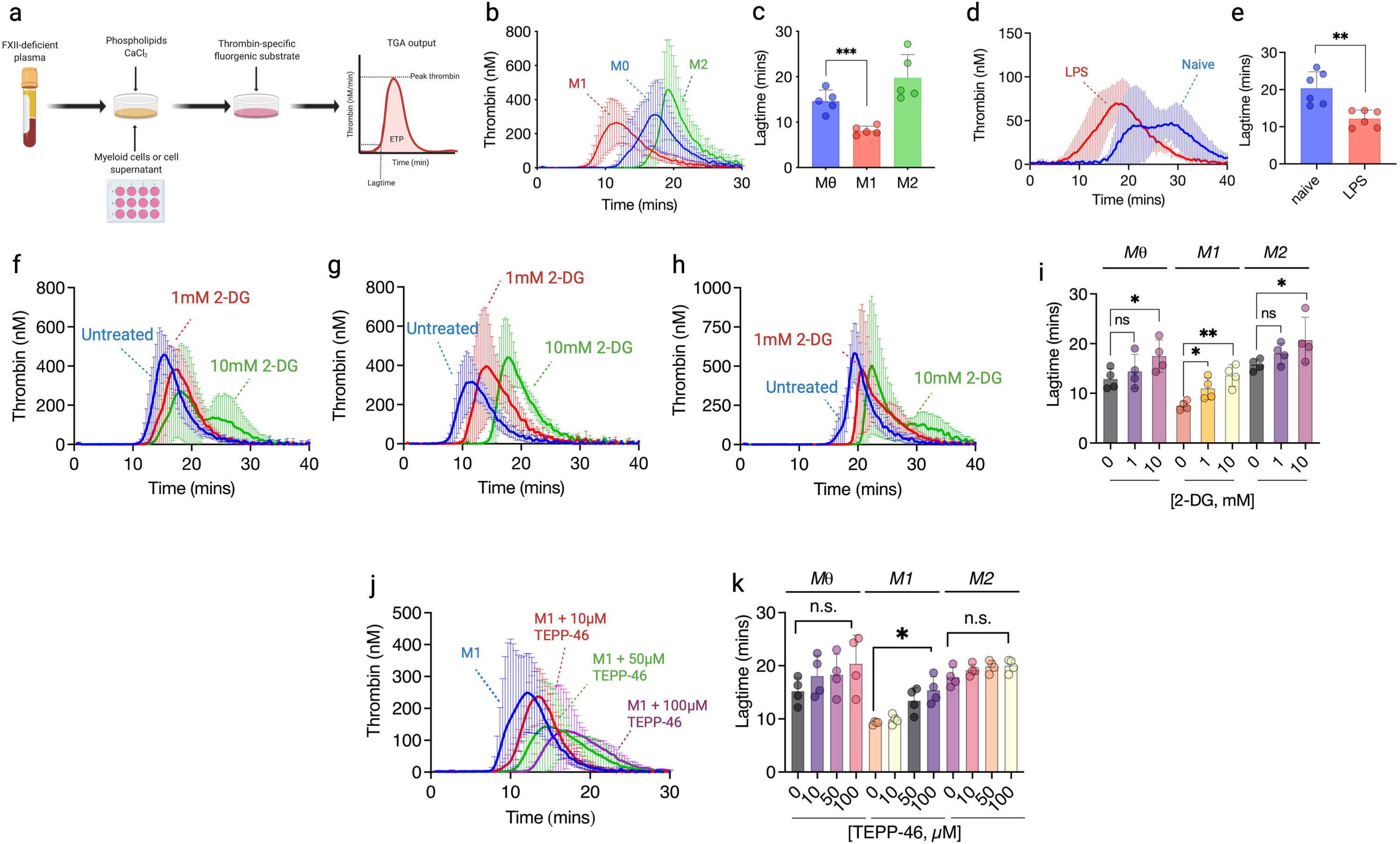
Glycolysis is essential for pro-inflammatory myeloid cell procoagulant activity. **(a)** Schematic diagram outlining myeloid cell-based CAT assay, in which incubated human monocytes or mouse bone marrow-derived macrophages (BMDMs) represent the sole source of TF activity. **(b)** Thrombograms for Mε, M1 (20 ng/mL IFNγ and 100 ng/mL LPS) and M2 (20 ng/mL IL-4 and IL-13) polarised BMDMs for 18 hr and **(c)** associated lag-times. **(d)** Thrombograms for CAT assay with human primary monocytes activated with 100ng/mL LPS for 18 hr and **(e)** lag-times. **(f)** Naïve, **(g)** M1 and **(h)** M2 BMDMs were pre-treated with 2-DG for 3 hrs (1-10mM), polarised, and CAT assays were performed, allowing **(i)** lag-times to be determined. **(j)** BMDMs were pre-treated with TEPP-46 (10-100µM) for 3 hrs and polarised before CAT assays were performed and **(k)** lag-times determined. A paired Student t-test or one-way ANOVA was used where appropriate to determine statistical significance with *P≤0.05, **P≤0.01 and ***P≤0.001 for 3-6 independent experiments measured in duplicate.

Induction of procoagulant activity was not limited to M1 polarisation. Notably, treatment with a TLR7/8 agonist, single-stranded RNA (ssRNA), that mimics ssRNA viral infection also yielded a qualitatively similar pattern of increased *PROCR* (9-fold), *F10* (15-fold) and *F3* (7-fold) expression to that of M1-polarised BMDMs **(Supplementary Figure 1e)**. We also observed reduced lagtime in both BMDMs **(Supplementary Figure 3a-b)** and human primary monocytes **(Supplementary Figure 3c-e)** treated with ssRNA. Viral infection has been reported to induce TF-containing microparticle release [38]. To assess the potential contribution of TF-containing extracellular vesicles, supernatants from treated cells were assessed. No difference in thrombin generation capacity between naive, M1 and M2 BMDM-derived supernatants, taken at several different time points, was observed **(Supplementary Figure 4a-h)**, Supernatants from ssRNA-treated BMDMs decreased lag-time, indicative of TF-containing extracellular vesicle release from ssRNA-treated BMDMs **(Supplementary Figure 4i-l)**.

Macrophage polarisation requires extensive metabolic reprogramming to achieve an M1 or M2 phenotype [24–26]. M1 polarisation and ssRNA treatment in BMDMs enhanced basal glycolysis, whereas M2 polarisation had minimal effect **(Supplementary Figure 5a)**. To evaluate the role of glycolytic reprogramming in fuelling myeloid cell-mediated procoagulant activities, murine and human myeloid cells were pre-treated with 2-deoxy-d-glucose (2-DG), a glucose analogue which inhibits glycolysis by competitive inhibition of hexokinase **(Supplementary Figure 5b)**. Notably, 2-DG treatment significantly prolonged CAT lag-time in naïve, M1 and M2 phenotypes in BMDMs **(Figure 1f-i)**, and was most effective in M1 BMDMs **(Figure 1g and 1i)**. No significant change in ETP or peak thrombin with 2-DG pre-treatment was observed. 2-DG treatment also caused lag time extension in LPS-stimulated human monocytes, M1-polarised HMDMs, ssRNA-treated BMDMs and human monocytes, restoring lag time to that observed in the presence of untreated cells **(Supplementary Figure 5)**. Cell viability following 2-DG treatment was examined by flow cytometry and found to be unaffected by glycolysis inhibition **(Supplementary Figure 6)**. These data highlight a direct role for glycolysis in enabling the procoagulant properties of pro-inflammatory monocytes and macrophages.

Small molecule PKM2 activators promote PKM2 tetramerisation, which prevents nuclear localisation and inhibits glycolysis by diverting pyruvate from the TCA cycle [39]. To investigate the effect of PKM2 tetramerisation on myeloid cell hypercoagulability, macrophages were pre-treated with PKM2 activator TEPP-46 and procoagulant activity was assessed by myeloid cell-based CAT assay. TEPP-46 treatment of M1 macrophages delayed lag-time by 5 minutes compared to untreated M1 macrophages **(Figure 1j)**. As anticipated, given that PKM2 tetramerisation occurs only in M1 macrophages, there was no change in lag-time for TEPP-46-treated naïve and M2 macrophages **(Figure 1j-k)**.

### Increased myeloid cell TF expression and decryption requires glycolytic reprogramming

To investigate the mechanisms through which 2-DG diminished procoagulant activity in activated myeloid cells, we next evaluated the role of glycolytic activity upon TF expression and activity in BMDMs. 2-DG caused an 8-fold and 10-fold reduction in TF cell membrane expression in both naïve and M1 polarised BMDMs, respectively **(Figure 2a)**. In parallel, gene expression analysis of 2-DG-treated M1 BMDMs showed a 14-fold reduction in *F3* gene expression levels compared to untreated M1 BMDMs **(Figure 2b)**. Moreover, an 11-fold attenuation in *F10* gene expression and a significant reduction in FX protein levels was also observed **(Figure 2c and d)**. To assess the effect of 2-DG on the expression of transcription factors known to regulate TF expression, BMDMs were treated with 2-DG and expression of EGR1 and AP-1 proteins (Jun and Fos sub-families) were measured. Notably, expression of EGR1 and AP-1 transcription factors were, as anticipated, significantly increased in M1-polarised BMDMs, however, their expression was significantly restricted by 2-DG treatment **(Figure 2e)**. We hypothesised that inhibition of glycolytic reprogramming might also impair the biosynthetic pathways necessary for membrane TF decryption. Membrane phosphatidylserine (PS) exposure was enhanced in M1 BMDMs compared to naïve BMDMs **(Figure 2f-g)** but was significantly suppressed by the presence of 2-DG. Recently, membrane translocation of acid sphingomyelinase (ASMase) has been shown to promote TF decryption via membrane sphingomyelinase degradation [40]. To determine whether ASMase membrane re-localisation is influenced by glycolysis, we visualised ASMase expression and membrane translocation in M1 polarised BMDMs in the presence or absence of 2-DG. M1 BMDM polarisation prompted formation of ASMase punctae proximal to the membrane surface **(Figure 2h-j)**. However, 2-DG attenuated ASMase punctae formation and diminished ASMase expression at the membrane surface **(Figure 2h-j)**.

**Figure 2:**
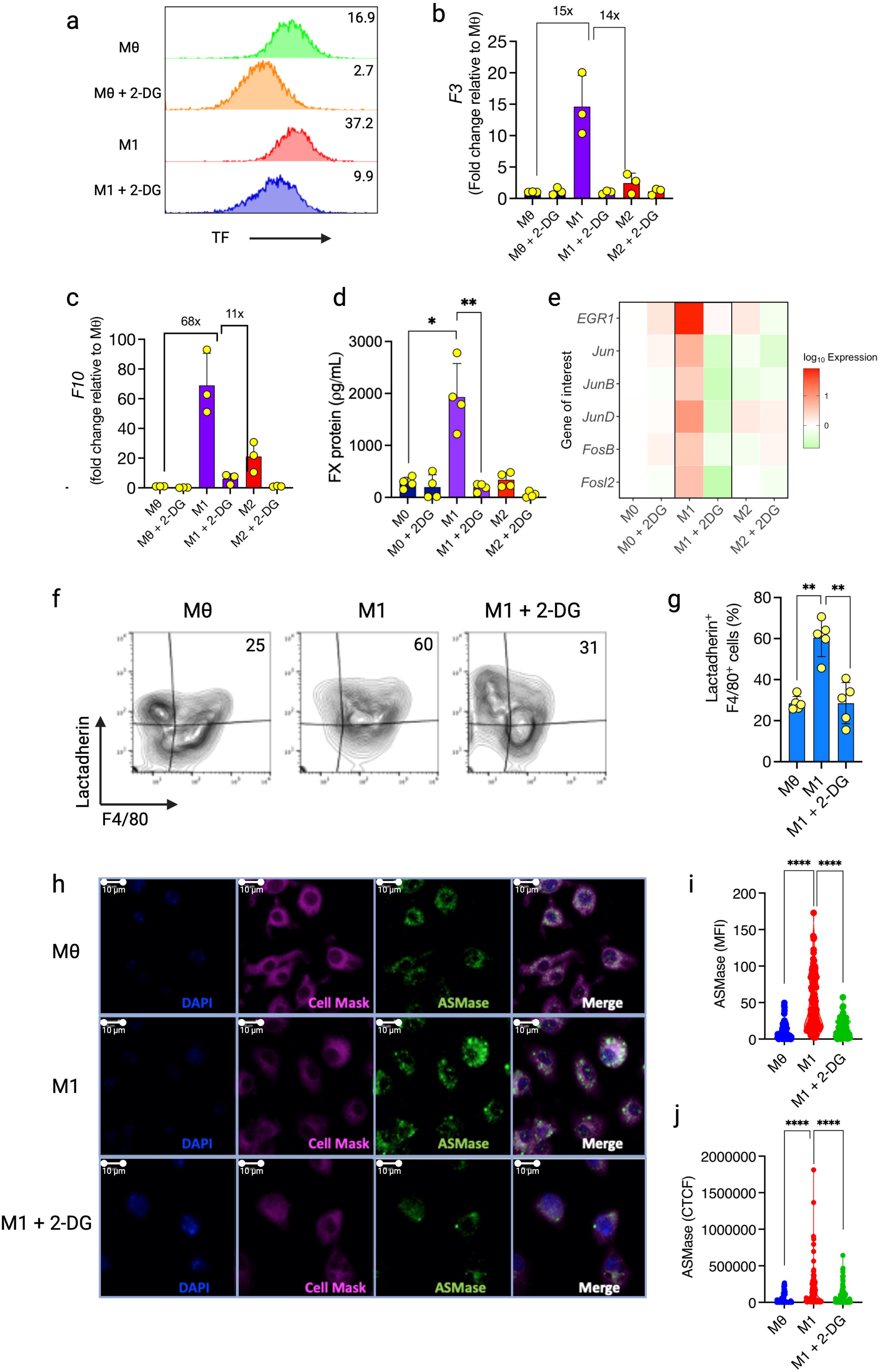
Increased myeloid cell TF expression and decryption require glycolytic reprogramming. **(a)** 2-DG-treated BMDMs either remained untreated or were M1 polarised before TF protein cell surface expression was assessed by flow cytometry. Representative histograms depicting TF^+^ cells (%) from 3 independent experiments are shown. **(b)** *F3* and **(c)** *F10* mRNA levels were determined by RT-qPCR. **(d)** Supernatant FX was determined by ELISA. **(e)** Heat map showing EGR1 and AP-1 transcription factors mRNA levels for Mε, M1 and M2 BMDMs with or without 2-DG pre-treatment were determined by RT-qPCR. **(f-g)** BMDM phosphatidylserine externalisation was measured using fluorescently-labelled lactadherin, and the percentage of lactadherin-bound cells was determined in 5 independent experiments**. (h)** Confocal microscopy was used to determine ASMase expression and localisation on BMDMs. DAPI (nucleus) and Cell Mask (cell membrane) were used to create masks for essential cellular organelles for image quantification. **(i)** Corrected total cell fluorescence (CTCF) and **(j)** mean fluorescence intensity for ASMase expression were calculated using Fiji software. A paired Student t-test, one-way ANOVA or Mann-Whitney U test was used where appropriate to determine statistical significance with *P≤0.05, **P≤0.01,****P≤0.0001 for 4-6 independent experiments measured in duplicate.

### Inhibition of glycolysis restricts myeloid cell PAI-1/2 secretion to enhance plasmin generation

Macrophages can enhance fibrinolysis via u-PA release to promote plasminogen activation [41–43]; however, inflammatory stimuli impairs this activity by inducing macrophage PAI-1 expression [43–45]. We first measured fibrinolysis-associated gene expression in polarised and ssRNA-treated BMDMs. As expected, significantly increased *SERPINE1* and *SERPINB2* gene expression, which encodes PAI-1 and PAI-2 respectively, was observed in M1 macrophages **(Figure 3a)**. In contrast, ssRNA did not affect *SERPINE1* expression, but *SERPINB2* expression was markedly increased **(Figure 3a)**. We used a plasma-free plasmin generation assay which allowed us to examine the ability of myeloid cells to facilitate tPA-dependent plasmin generation and capture cell-derived PAI-1/2 regulation of plasmin generation activity. t-PA mediated plasmin generation was ablated in the presence of M1 polarised BMDMs compared to naïve BMDMs **(Figure 3b)**. In contrast, M2-polarised BMDMs exhibited significantly increased plasmin generation compared to naïve macrophages **(Figure 3b)**.

**Figure 3:**
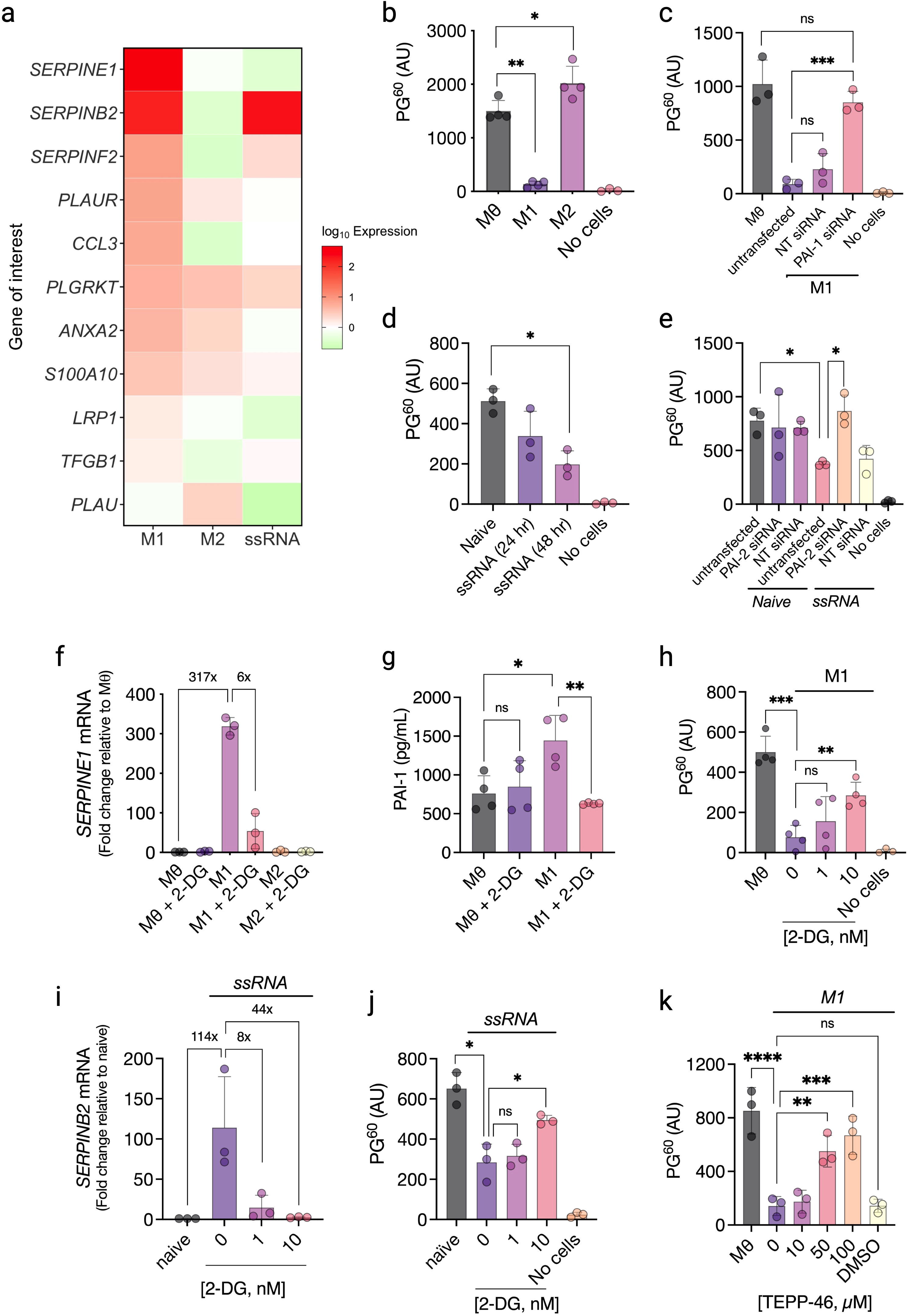
Inhibition of glucose metabolism restricts myeloid cell PAI-1/2 secretion to enhance plasmin generation. 2-DG-treated BMDMs were polarised towards M1 or M2 or treated with ssRNA before gene expression and plasmin generation analysis. **(a)** Heat-map showing fibrinolysis-associated gene expression profiles for M1, M2, and ssRNA-treated BMDMs. **(b)** t-PA-mediated plasmin generation in the presence of polarised BMDMs was measured using a plasmin-specific fluorogenic substrate, and the fluorescence reading after 60 mins (PG^60^) was determined. **(c)** BMDMs were transfected with *SERPINE1* siRNA or non-targeting (NT) siRNA (both 20nM), before M1 polarisation and plasmin generation analysis. **(d)** Plasmin generation analysis was performed with ssRNA-treated BMDMs, as previously described. **(e)** BMDMs were transfected with 20nM *SERPINB2* siRNA or non-targeting (NT) siRNA before ssRNA exposure, and plasmin generation was assessed. **(f)** 2-DG-treated BMDMs were polarised before assessing *SERPINE1* mRNA levels, **(g)** PAI-1 production by ELISA, and **(h)** plasmin generation. **(i)** 2-DG-treated BMDMs were treated with ssRNA before measuring *SERPINB2* gene expression and **(j)** plasmin generation. **(k)** Polarised BMDMs were treated with TEPP-46 (10-100µM), then M1 polarised, and plasmin generation was determined. A paired Student t-test or one-way ANOVA was used where appropriate to determine statistical significance with *P≤0.05, **P≤0.01, ***P≤0.001, ****P≤0.0001 for 3-4 independent experiments measured in duplicate.

To confirm the role of PAI-1-dependent, M1 macrophage-mediated inhibition of plasmin generation, *SERPINE1* gene silencing using targeted siRNA was performed. siRNA knockdown efficacy was confirmed via *SERPINE1* mRNA expression by qPCR, which showed a 490-fold decrease upon PAI-1 siRNA transfection compared to untransfected M1 BMDMs **(Supplementary Figure 7a)**. Untransfected M1 BMDMs significantly inhibited plasmin generation and no significant change in plasmin generation was observed in BMDMs treated with non-targeting siRNA **(Figure 3c)**. However, *SERPINE1* siRNA-treated M1 BMDMs exhibited significantly increased t-PA-mediated plasmin generation, comparable to that observed in the presence of untreated BMDMs **(Figure 3c)**, confirming the specific role of inflammation-induced macrophage PAI-1 release in the inhibition of t-PA mediated plasmin generation.

As in M1 BMDMs, ssRNA treatment caused attenuation of t-PA-mediated plasmin generation. Reduced plasmin generation was observed as ssRNA exposure time increased **(Figure 3d)**. Interestingly, ssRNA-treated BMDMs exhibited significantly increased *SERPINB2* expression (encoding PAI-2), which was increased 200-fold relative to *SERPINB2* expression in naïve BMDMs **(Figure 3a)**. To establish whether the observed inhibition of plasmin generation in ssRNA-treated macrophages arose because of increased PAI-2 expression, *SERPINB2* gene silencing in ssRNA-treated BMDMs was performed using *SERPINB2*-targeted siRNA **(Supplementary Figure 7b)**. t-PA-mediated plasmin generation capacity in *SERPINB2*-silenced ssRNA-treated BMDMs was double that observed in naïve BMDMs **(Figure 3e)**. This data demonstrates a novel role for PAI-2 extracellular release in regulating ssRNA-exposed macrophage fibrinolytic activity.

We next sought to establish whether re-programming cellular metabolism was important in enabling pro-inflammatory macrophage anti-fibrinolytic activity. *SERPINE1* gene expression was significantly reduced in M1 polarised BMDMs following 2-DG treatment **(Figure 3f)**. This led to a ∼50% reduction in PAI-1 secretion from 2-DG-treated M1 BMDMs compared to untreated M1 BMDMs **(Figure 3g)**. 2-DG treatment reversed M1 macrophage antifibrinolytic activity and increased M1 macrophage-dependent, t-PA-mediated plasmin generation by 4-fold **(Figure 3h)**. There was no change in t-PA-mediated plasmin generation upon 2-DG pre-treatment in naive and M2 cell populations. 2-DG treatment in ssRNA-treated BMDMs mediated a 100-fold decrease in *SERPINB2* expression **(Figure 3i)** which resulted in restored macrophage fibrinolytic activity, doubling t-PA-mediated plasmin generation with ssRNA-treated BMDMs present **(Figure 3j)**. To investigate the effect of PKM2 tetramerisation on myeloid cell antifibrinolytic activity, BMDMs were pre-treated with PKM2 activator TEPP-46 and plasmin generation was assessed. As with 2-DG, we observed a dose-dependent increase in t-PA-mediated plasmin generation in the presence of TEPP-46 **(Figure 3k)**. Together, this data indicates that PAI-1 and PAI-2 expression mediate macrophage anti-fibrinolytic activity in response to different pro-inflammatory stimuli, which can be ameliorated using pharmacological inhibitors of glucose metabolism.

### Myeloid cell glycolytic reprogramming increases EPCR expression and accelerates APC generation

Endogenous anticoagulant pathways are typically down-regulated on endothelial cells in response to pathogen- or cytokine stimulation [46–48]. Macrophage polarisation had no effect on tissue factor pathway inhibitor (TFPI) or antithrombin expression **(Supplementary Figure 1e)**, however, *PROCR* expression in M1 BMDMs was increased 250-fold compared to naïve or M2 BMDMs **(Figure 4a)**. As previously reported[49], EPCR cell surface expression was largely absent on naïve BMDMs, but expression was almost completely restored by M1 polarisation **(Figure 4b-c)**. In contrast, TM expression was not significantly affected by M1 polarisation, but was enhanced 3-fold on M2 macrophages **(Figure 4d-e)**. In polarised HMDMs, similar changes in *PROCR* and *THBD* gene activity and EPCR and TM surface expression were also observed (**Supplementary Figure 8**). Both naïve, M1 and M2 BMDMs supported protein C activation, consistent with TM expression in each macrophage sub-type **(Figure 4f)**. M1-polarised BMDMs, however, exhibited ∼3-fold enhanced APC generation compared to naïve and M2-polarised BMDMs, consistent with EPCR expression being uniquely up-regulated in this macrophage subset.

**Figure 4:**
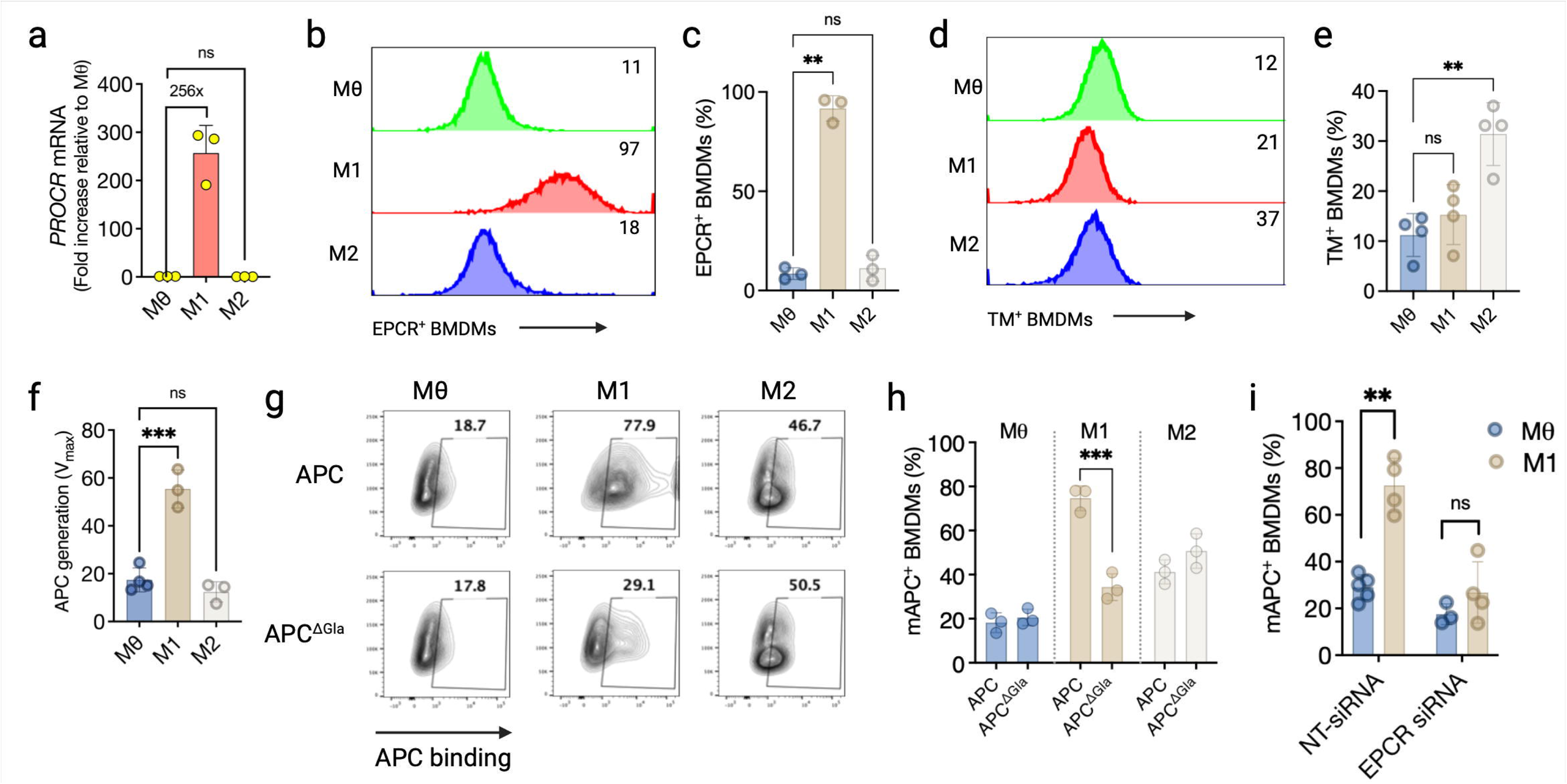
Myeloid cell glycolytic reprogramming increases EPCR expression and accelerates APC generation. **(a)** *PROCR* expression in polarised BMDMs was determined by RT-qPCR. BMDMs were polarised, and surface expression of **(b-c)** EPCR and **(d-e)** TM was determined by flow cytometry. Results presented are representative histograms depicting the % of live EPCR^+^ or TM^+^ cells. **(f)** Murine protein C activation by thrombin in the presence of untreated or polarised BMDMs was measured by incubating cells with 100nM mouse PC and 5nM thrombin for 1hr at 37°C. Hirudin was then added to stop the reaction and 50µL of APC chromogenic substrate was added to detect PC activation. **(g-h)** Polarised BMDMs were incubated with 50nM fluorescently-labelled wild-type APC or Gla-domainless APC (APC^ΔGla^) for 2 hours before flow cytometry measurement of cell binding. **(i)** BMDMs were transfected with 20nM *PROCR* siRNA or non-targeting (NT) siRNA for 24 hours before M1 polarisation. The following day, BMDMs were incubated with 50nM fluorescently-labelled APC for 2 hours prior to allow detection of cell binding by flow cytometry. A paired Student t-test or one-way ANOVA was used where appropriate to determine statistical significance with *P≤0.05, **P≤0.01, ***P≤0.00 for 3-7 independent experiments measured in duplicate.

Fluorescently-labelled APC binding was observed on naïve macrophages, but was significantly enhanced (∼4-fold) in the presence of M1-polarised macrophages **(Figure 4g-h)**. To ascertain whether this increased binding on M1 macrophages was due to EPCR binding, we measured macrophage binding of an APC variant truncated to remove the N-terminal Gla domain (APC^ΔGla^) required for EPCR [50], but not Mac-1 binding[49]. Whereas APC^ΔGla^ binding to naïve and M2-polarised BMDMs was like that observed in the presence of wild-type APC, binding to M1-polarised BMDMs was strikingly reduced, suggesting M1 polarisation-induced EPCR expression was critical for enhanced APC binding. Intriguingly, we also observed a 2-3-fold increase in wild-type APC and APC^ΔGla^ binding to M2-polarised BMDMs independent of EPCR expression **(Figure 4g-h)**. siRNA-mediated EPCR knockdown in M1-polarised BMDMs also significantly reduced APC binding to that observed when APC was incubated with naïve macrophages, underscoring the requirement for EPCR in mediating enhanced APC binding to macrophages after exposure to pro-inflammatory stimuli **(Figure 4i)**.

### Tissue-resident and peripheral TF^+^ and EPCR^+^ myeloid cells are present in mice with immunometabolic disease

Based on this data, we hypothesised that EPCR^+^ tissue-resident myeloid cells would also arise predominantly within inflamed tissue *in vivo*. To test this, adipose tissue macrophages (ATMs) derived from either normal control diet (CD)-fed mice or obese mice fed a high-fat diet (HFD) were assessed **(Figure 5a)**. ATMs exhibit chronic metabolic dysfunction, procoagulant activity and anti-fibrinolytic properties [51,52], but their ability to regulate anticoagulant pathways remains unknown. Using an existing transcriptomic dataset [53] of ATMs isolated from CD and HFD-fed mice, we found that ATMs from HFD-fed mice exhibit an “M1-like” phenotype, with several established canonical M1 genetic markers (*tnf*, *Il1b*, *Nos2*, *CD80*) significantly upregulated compared to mice fed a normal diet **(Figure 5b and Supplementary Table 2)**, which is in keeping with previous literature [51,53,54]. Additionally, both *F10* and *SERPINE1* expression was significantly increased in ATMs from HFD-fed mice **(Figure 5b and Supplementary Table 2)**. Accordingly, we observed a significant reduction in plasma CAT lag-time on ATMs isolated from HFD-fed obese mice compared to CD-fed mice **(Figure 5c-d)**. Moreover, plasmin generation in the presence of ATMs derived from HFD-fed mice was significantly reduced compared to that observed in ATMs from CD-fed mice **(Figure 5e)** confirming previous findings of the anti-fibrinolytic activity of ATMs from obese mice [52,55]. We next measured protein C pathway receptor expression and APC generation on isolated ATMs from obese and control mice. Interestingly, EPCR^+^ ATMs were 8-fold more prevalent in adipose tissue isolated from HFD-fed mice compared to ATMs isolated from CD-fed mice **(Figure 5f-g)**. There was also a 10-fold increase in ATMs expressing TM in HFD-fed mice **(Figure 5h-i)** (See **Supplementary Figure 9** for macrophage characterisation and gating strategies). Consistent with increased TM and EPCR expression, 10-fold more APC was generated in the presence of ATMs from HFD-fed mice compared to CD-fed mice **(Figure 5j)**. These data demonstrate that the procoagulant and antifibrinolytic activity of tissue-resident adipose tissue macrophages from obese mice is accompanied by increased protein C pathway receptor expression and APC generation capacity.

**Figure 5:**
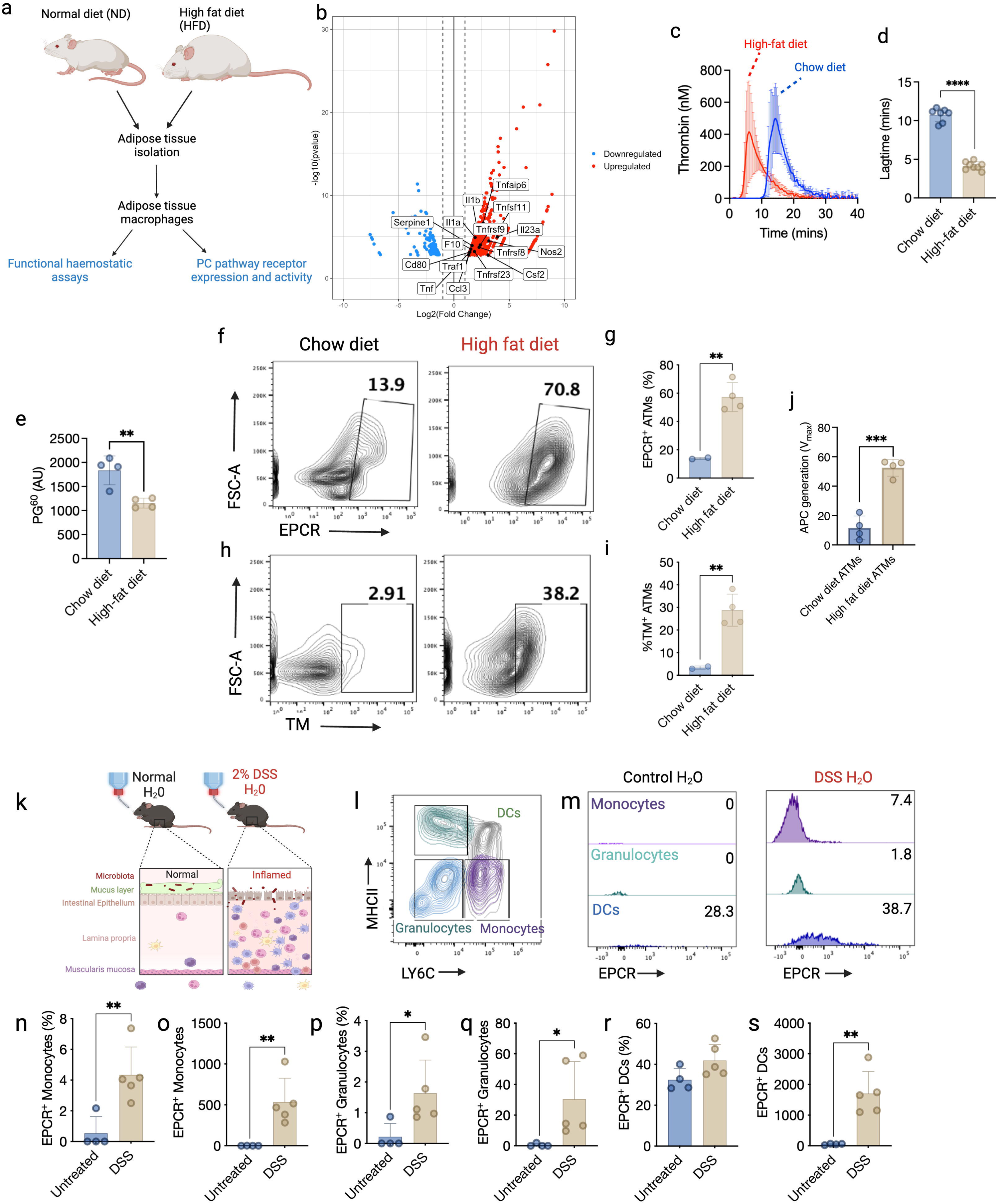
Tissue-resident and peripheral infiltrating TF+ and EPCR^+^ myeloid cells are present in mice with immunometabolic disease. **(a)** Schematic representation of experimental plan for high fat diet (HFD)-induced obesity model. Mice were fed a control chow diet (CD, n=7) or HFD (n=8) for 12 weeks before being sacrificed, and WAT and ATMs were collected. **(b)** Volcano plot of differential expression of genes between adipose tissue macrophages (ATMs) isolated from control diet and high-fat diet fed mice with canonical M1 markers highlighted (analysis carried out on GSE158627 dataset). **(c)** Myeloid cell-based CAT assays were performed in the presence of ATMs isolated from CD and HFD-fed mice, enabling **(d)** lag-times to be determined. **(e)** plasmin generation in the presence of ATMs isolated from HFD and CD mice was determined. **(f-g)** EPCR cell surface expression and **(h-i)** TM cell surface expression was determined in CD and HFD ATMs. **(j)** Protein C activation in the presence of CD and HFD ATMs was also performed. **(k)** Schematic description of the DSS-induced colitis model. Mice were supplemented with 2% DSS in their drinking water for 5 consecutive days to induce colitis (untreated n=4, DSS n=5) **(l)** Following sacrifice, the colons of the mice were harvested, digested, and analysed for surface expression of leukocyte population markers by flow cytometry. **(m)** EPCR expression and the total number of EPCR-expressing cells were then measured in the individual monocytes **(n-o)**, granulocytes **(p-q)**, and DC populations **(r-s)**. An unpaired Student t-test was used to determine statistical significance with *P≤0.05, **P≤0.01, ***P≤0.001 and ****P≤0.0001 measured in duplicate.

We next characterised EPCR expression on innate immune cells isolated from intestinal tissue of colitic or untreated mice. **(Figure 5k)**. Colitis is characterised as a chronic inflammatory condition in which “M1-like” inflammatory macrophages are a major driver of disease pathophysiology [56,57] At disease peak, colons were harvested, and leukocytes were isolated from the lamina propria. The unfractionated leukocyte population was then stained for EPCR and specific cell surface markers to enable identification of the different innate immune cell subtypes present, including monocytes (CD45^+^CD3^−^CD11b^+^MHCII^low^LY6C^hi^), dendritic cells (CD45^+^CD3^−^ CD11b^+^MHCII^high^LY6C^low^) and granulocytes (CD45^+^CD3^−^CD11b^+^MHCII^low^LY6C^low^; **Figure 5l)**. Interestingly, enhanced EPCR expression was observed in all innate cell subsets isolated from inflamed intestinal tissue of DSS-treated mice compared to control mice **(Figure 5m)** and a significant increase in EPCR^+^ monocyte, granulocyte and DCs infiltrating the colon at disease peak was observed **(Figure 5n-s and Supplementary Figure 10**).

## DISCUSSION

Previous studies have highlighted the capacity of both tissue-resident macrophages in inflamed tissue and circulating monocytes to promote coagulation via enhanced TF activity [1,58,59], although the mechanisms enabling this transition to a procoagulant phenotype are not fully understood. Myeloid cells exhibit phenotypic plasticity shaped by their microenvironment, tissue location and pathogen exposure [60]. In this study, we used *ex vivo* cell-based coagulation assays and *in vivo* inflammatory disease models to show how induction of myeloid cell metabolic reprogramming in response to inflammatory stressors modulates plasma procoagulant, antifibrinolytic and anticoagulant activities. M1 macrophage polarisation promoted TF decryption or increased procoagulant activity, whereas M2 macrophages exhibited limited TF-dependent procoagulant activity, in keeping with the procoagulant activity of myeloid cells in acute inflammatory disease models [3,40,61]. Similarly, tissue-resident macrophages in inflamed tissues exhibit potent anti-fibrinolytic activity via up-regulated PAI-1 secretion [55]. In our study, macrophage polarisation toward either M1 or M2 phenotypes provoked divergent effects on macrophage fibrinolytic activity. The ability of naive and M2 polarised macrophages to drive plasmin generation is likely due to u-PA expression by monocytes and macrophages [41,62,63], with one study linking u-PA over-expression to an M2 macrophage phenotype in a murine fibrosis model [63]. These data suggest a delicate balance between macrophage plasminogen activators and inhibitors to regulate local plasmin activity around an injury site, in which the initial pro-inflammatory response to tissue injury is accompanied by inhibition of local fibrinolytic activity to enable stable clot formation. Although our data is in keeping with the procoagulant and anti-fibrinolytic activity of pro-inflammatory myeloid cells in acute inflammatory disease models [40,61] and clinical studies [64,65], they contrast with a recent study that found human monocyte-derived M2-polarised macrophages exhibited enhanced TF expression and increased release of TF-containing extracellular vesicles, with no change in TF activity in human M1-polarised macrophages [66]. The reasons for these divergent findings are currently unknown, but may arise due to differences in culture conditions during macrophage differentiation, or due to differences in cytokine concentration and/or exposure time during polarisation.

To explore whether different pro-inflammatory stressors provoked similar or distinct alterations in myeloid cell haemostatic activity, we also evaluated thrombin and plasmin generation in the presence of ssRNA-exposed myeloid cells that mimic ssRNA virus infection. Viral infections are commonly associated with haemostatic dysregulation [67] and ssRNA viruses such as Human Immunodeficiency Virus (HIV), Influenza A and SARS-CoV-2 (which act as potent agonists of TLR7 in mice and TLR8 in humans)[68] are associated with increased risk of thromboembolic events in infected patients [69–73]. In HIV patients, circulating TF^+^ monocytes are increased compared to uninfected individuals and persist even after viral suppression [74]. Furthermore, TF^+^ monocytes contribute to the coagulopathy observed in a simian immunodeficiency virus (SIV) primate infection model [70]. Plasma microvesicle TF activity is significantly associated with mortality in ICU-admitted influenza patients [75] and SARS-CoV-2 infection causes increased monocyte TF expression and elevated plasma TF^+^ extracellular vesicle levels [38,76]. Consistent with these studies, human peripheral monocytes and murine macrophages became more procoagulant after TLR7/8 activation when evaluated in our myeloid cell-based CAT assay. In contrast to polarised M1 macrophages, PAI-2 was required for TLR7 agonist-mediated macrophage anti-fibrinolytic activity, highlighting a novel role for macrophage PAI-2 in response to viral infection. Unlike PAI-1, PAI-2 is thought to be mostly contained intracellularly in leukocytes, with only a small amount typically secreted into the extracellular space [77,78]. Despite this, PAI-2 can play an important role in the innate immune response to pathogens. Several studies have shown that PAI-2 can inhibit myeloid cell apoptosis [79], modulate monocyte proliferation and differentiation [80] and may be protective against viral infections such as hepatitis A [81]. PAI-2 has also been shown to impact thrombus resolution *in vivo,* as PAI-2 deficiency causes enhanced thrombus lysis in a murine DVT model [82].

Myeloid cell metabolic dysregulation is observed in many inflammatory disease states [83,84]. The transition of macrophages from naïve to a pro-inflammatory phenotype is typically accompanied by a shift in cellular metabolism from oxidative phosphorylation as the principal means of energy generation to glycolysis [24]. Previous studies have demonstrated that inhibition of glycolysis with 2-DG suppresses LPS-induced cytokine release from murine macrophages, dependent upon succinate accumulation to stabilise HIF1α and promote pro-inflammatory signalling [25]. We observed that myeloid cell glycolytic metabolism inhibition provoked a hypocoagulable myeloid cell phenotype caused by diminished cell surface TF activity and enhanced fibrinolytic function. 2-DG inhibited TF-dependent procoagulant activity in LPS- and ssRNA-treated macrophages and monocytes, consistent with a novel role for glycolytic metabolism in fuelling TF expression and decryption. Glycolytic metabolism was also necessary for the antifibrinolytic activity of M1 polarised and ssRNA-treated macrophages, which was mediated by enhanced PAI-1/2 expression and release. It is currently unknown how glycolysis controls macrophage PAI-1/2 expression, however, several studies have shown that pyruvate, the end product of glycolysis, upregulates PAI-1 expression through a HIF-1α dependant pathway [85–87]. Both TF and PAI-1 expression share common transcriptional upregulation via engagement of transcriptional factors such as EGR1, AP-1 and NF-κ;B [88–91]. M1 BMDMs showed increased EGR1 and AP-1 transcription factor expression, which was significantly attenuated by 2-DG treatment. This suggests that inhibition of glycolysis by 2-DG regulates procoagulant and antifibrinolytic gene expression via control of shared upstream transcriptional elements. Furthermore, it remains possible that changes in TF, FX and PAI-1 expression may also be influenced by cytokine release secondary to M1 polarisation. Further studies are ongoing to disentangle the complex mechanisms underlying glycolytic control of myeloid cell procoagulant activity. We also observed hypercoagulability and hypofibrinolytic activity in adipose tissue macrophages isolated from mice fed a high-fat diet, further supporting a key role for enhanced glycolytic activity in driving immunothrombotic activity *in vivo*. These data suggest pharmacological strategies to promote metabolic re-wiring of activated monocytes [92] could effectively regulate inflammation-mediated procoagulant activity.

In contrast to its established role in mediating anti-inflammatory cell signalling [49,93,94] the role of myeloid cell type, polarisation, or *in vivo* inflammatory status on APC generation and anticoagulant activity is unknown. We confirmed that on human macrophages and murine bone marrow-derived macrophages [49], EPCR expression is normally limited, but rapidly up-regulated by exposure to pro-inflammatory stimuli. Surprisingly, EPCR expression was markedly enhanced on bone marrow- and tissue-derived myeloid cells in response to any pro-inflammatory stimuli tested in this study, suggesting myeloid EPCR is a sensitive marker of local inflammatory activity. Additionally, inflamed myeloid cells supported enhanced APC generation, but our myeloid cell CAT data suggest this was insufficient to counterbalance the increased procoagulant activity caused by increased myeloid TF expression and activity. However, M1 macrophages exhibited consistently reduced peak thrombin compared to naïve and M2 macrophages, consistent with enhanced APC generation. Instead, increased APC generation on stimulated myeloid cells may be particularly important for enhanced APC-mediated anti-inflammatory activity, mediating a similar regulatory role as the cytokine interleukin-10, which is also up-regulated and secreted in response to pro-inflammatory challenge [95]. Further studies are underway to characterise the interaction of glucose metabolism, EPCR expression and its role in the hypercoagulability of pro-inflammatory monocyte subsets isolated from patient cohorts with inflammatory disease and increased VTE risk.

In summary, we demonstrate that metabolic reprogramming arising from macrophage activation is essential for enhanced TF activity and diminished fibrinolytic activity, and has divergent impacts on APC-dependent anticoagulant and anti-inflammatory activities. As such, pharmacological approaches to modulate innate immune cell metabolic dysregulation may represent a novel strategy to mitigate VTE in at-risk patients.

## AUTHOR CONTRIBUTIONS

A.M.R and R.J.S.P devised the study, designed the experimental strategy, and wrote the manuscript. A.M.R, G.L., S.M., P.K., I.S., Y.H-S., S.A., T.R., T.R., A.M.C., S.H., J.S.O’D and P.T.W. performed experiments and analyzed data. All authors reviewed and contributed to the final version of the manuscript.

## ACKNOWLEDGEMENTS

Grant support for R.J.S.P. is provided by Science Foundation Ireland (21/FFP-A/8859), The National Children’s Research Centre (C/18/3) and Health Research Board (ILP-POR-2022-060).

