## Supplementary Materials for "Glycolytic reprogramming fuels myeloid cell-driven hypercoagulability"

### SUPPLEMENTARY MATERIALS AND METHODS

#### *Generation of bone marrow-derived macrophages*

Female wild-type C57/BL6 mice aged between 6-12 weeks were euthanised by cervical dislocation, and hind legs dissected, removed, and stored in high glucose DMEM (Merck) supplemented with 10% foetal bovine serum (ThermoFisher Scientific), 1U/mL penicillin and 0.1mg/mL streptomycin (Merck). Macrophages were isolated as previously described [1,2]. Isolated macrophages were re-suspended in supplemented DMEM containing 20% M-CSF derived from L929 cells and incubated for six days, receiving 1mL M-CSF and 4mL fresh growth media on day 3. After six days, macrophages were detached using PBS containing 5mM EDTA (Merck) and plated for experiments in complete DMEM containing 10% M-CSF.

#### *Culture of human macrophages*

PBMCs were isolated from buffy coats by layering 20mL blood over 15 mL histopaque-1077 gradient (Sigma) and centrifuging at 400xg for 25 minutes with acceleration and brake set to 1. CD14+ monocytes were isolated using CD14+ microbeads positive selection (Miltenyi Biotec) and plated at 2.5 x $10^5$ cells/mL in RPMI containing 1U/mL penicillin and 0.1mg/mL streptomycin, and 10% human serum. Cells were left for 10 days to allow them to differentiate into macrophages. Where M-CSF was used for differentiation, recombinant M-CSF (Peprotech) was added to culture media (50 ng/mL). Nonadherent cells were removed on days four and seven, and media was replaced.

#### *Culture and differentiation of THP1 macrophages*

THP1 monocytic cell line were cultured in suspension in non-adherent flasks at a density of 1x$10^5$ cells per mL to 1x$10^6$ cells/mL in RPMI supplemented with 10% foetal bovine serum, 1U/mL penicillin, 0.1mg/mL streptomycin and 2 mM L-glutamine (Merck). To differentiate cells into adherent macrophages, cells were resuspended at a density of 2x$10^5$ cells/mL and cultured with 100nM para-Methoxyamphetamin (PMA, Merck) for 3 days. After this time, PMA was removed and new growth media added. Cells were allowed to rest for 1 day before polarisation towards M1 and M2 phenotypes with same cytokine concentrations as for primary human macrophages.

***PAI-1 ELISA***

PAI-1 levels released by treated macrophages were determined using DuoSet ELISA (R&D Systems) per the manufacturer's protocol.

***Murine FX ELISA***

FX in BMDM supernatants was assessed using Mouse FX ELISA Kit (FineTest, Quadratech) as per the manufacturer's protocol.

***Cellular glycolytic rate assay***

Real-time cell metabolic analysis was performed using the Seahorse XF Glycolytic Rate Assay kit (Agilent). $5 \times 10^4$ cells per well were seeded onto a specialised Seahorse XF96 cell culture microplate and left to adhere overnight. Cells were treated with 5µg/mL ssRNA/LyoVec (Invivogen) or 100ng/mL LPS for 3-6 hours. Supernatants were removed, and cells were washed with pre-warmed assay medium (Seahorse XF DMEM supplemented with 1mM pyruvate, 2mM glutamine and 10mM glucose). Cells were then incubated with 200µL/well assay medium in a non-$CO_2$ incubator for 45-60 minutes. Following this, the assay medium was again removed from the cells, and 180µL/well fresh assay medium was added. 20µL Rot/AA was loaded into port A, and 22µL 2-DG was loaded into port B. Cellular glycolysis was then determined, and basal glycolysis defined as the last glycoPER measurement before inhibitor injection. Once the glycolytic rate assay was completed, data were normalised to total protein per well, as determined by the BCA reagent. Alterations in cellular metabolism were determined by changes in OCR and ECAR using Wave Software 2.0.

***Macrophage siRNA transfection***

Cells were seeded at $2.5 \times 10^5$ cells per well on 24-well plates. On the day of transfection, media was removed and replaced with 400µL of OptiMEM (Gibco) containing 5% M-CSF. siRNAs were transfected into cells at a concentration of 20nM using Lipofectamine RNAiMax (Invitrogen). Untransfected controls containing OptiMEM were also included in each experiment. The following day, media was removed and replaced with DMEM containing 10% M-CSF derived from L-929 cells. 24 hours following transfection, cells were treated as required for downstream assays.

***Mouse lamina propria leukocyte isolation***

At experiment termination, the colons of untreated and DSS-treated mice were excised and opened longitudinally. Faecal contents were removed, washed, and colons were cut into 0.5-1cm segments and transferred to a 50ml conical tube containing pre-digestion buffer (HBSS (Merck) supplemented with 5mM EDTA and 5% FBS. Colon segments were then shaken horizontally at 37°C for 20 minutes at 200rpm and then washed in HBSS to remove residual EDTA. The segments were then cut into 1mm pieces and transferred to a 50-ml conical tube containing digestion buffer (HBSS supplemented with 1.6mg/ml collagenase D, 40µg/ml DNase and 5% FBS) and shaken horizontally at 37°C for 45 minutes at 200rpm. Complete dissociation of remaining intestinal tissue was achieved by vortexing the tubes for 10 seconds after digestion. All intestinal content was then passed through a 100µm cell strainer, and the resulting cell suspensions were pelleted by centrifugation at 2500rpm. The isolated lamina propria leukocytes were subsequently analysed by flow cytometry.

***Mouse lamina propria leukocyte flow cytometry analysis***

Following lamina propria leukocytes isolation all cells were fluorescently stained for EPCR expression and cell surface expression of leukocyte lineage markers. All fluorescently labelled antibodies used were purchased from eBiosciences™ (anti-EPCR, anti-CD45, anti-MHC-II, anti-LY6C, anti-CD11b, and anti-CD11c). Cell viability was first measured using the LIVE/DEAD™ Fixable Aqua Dead Cell Stain Kit (ThermoFisher Scientific), and cell numbers were determined using CountBright™ Absolute Counting Beads (ThermoFisher Scientific). Before staining, cells were first incubated with an anti-CD16/CD32 Monoclonal Antibody to block FC receptors (Invitrogen). Cells were then washed using PBS and incubated with LIVE/DEAD™ dye for 30 minutes at 4$^{o}$C. Cells were washed using PBS supplemented with 2% FBS and incubated with fluorescently labelled antibodies for 1 hour at 4$^{o}$C. Multi-parameter analysis was then performed on an LSR/Fortessa (BD) and analysed using FlowJo 10 software. Positive staining was assessed using fluorescence minus one (FMO) controls. Monocytes were classed as being CD45$^{+}$CD3$^{-}$CD11b$^{+}$MHCII$^{low}$LY6C$^{hi}$ [3][4], dendritic cells were classed as being CD45$^{+}$CD3$^{-}$CD11b$^{+}$MHCII$^{high}$LY6C$^{low}$ [5][6][4], and granulocytes were classed as being CD45$^{+}$CD3$^{-}$CD11b$^{+}$MHCII$^{low}$LY6C$^{low}$ [3][7].

To analyse the percentage of cells expressing EPCR the following gating strategy was used: FSC-A vs SSC-A → Live Cells (SSC-A vs LIVE/DEAD™ dye$^{-}$) → Single Cells

(FSC-A vs FSC – W) → Innate Leukocytes (LIVE/DEAD™ dye$^{-}$CD45$^{+}$CD3$^{-}$) → CD11b+ Innate Leukocytes (CD45$^{+}$CD11b$^{+}$) → Monocytes (MHCII$^{low}$LY6C$^{hi}$), DCs (MHCII$^{high}$ LY6C$^{low}$), granulocytes (MHCII$^{low}$LY6C$^{low}$) → EPCR$^{+}$. To analyse the total number of cells expressing EPCR CountBright™ Absolute Counting Beads (ThermoFisher Scientific) were used using the following strategy: Total Live Cells (SSC-A vs LIVE/DEAD™ dye$^{-}$) → Total Single Cells (FSC-A vs FSC – W) → Total Innate Leukocytes (LIVE/DEAD™ dye$^{-}$CD45$^{+}$CD3$^{-}$) → Total CD11b+ Innate Leukocytes (CD45$^{+}$CD11b$^{+}$) → Total Monocytes (MHCII$^{low}$LY6C$^{hi}$), Total DCs (MHCII$^{high}$ LY6C$^{low}$), Total Granulocytes (MHCII$^{low}$LY6C$^{low}$) → Total EPCR+ Monocytes (EPCR$^{+}$MHCII$^{low}$LY6C$^{hi}$), Total EPCR$^{+}$ DCs (EPCR$^{+}$MHCII$^{high}$ LY6C$^{low}$) and Total EPCR$^{+}$ Granulocytes (EPCR$^{+}$MHCII$^{low}$LY6C$^{low}$).

***Isolation and culture of adipose tissue macrophages***

When HFD-fed mice reached a weight of ~45g, they were sacrificed as described, and the white adipose tissue (WAT) from inguinal tissue was collected. WATs were placed in 6-well dishes with PBS supplemented with 1U/mL penicillin and 0.1mg/mL streptomycin. WAT was minced using scissors and then transferred to 50mL tubes. Collagenase digestion was used to obtain the stromal vascular fraction from the WAT. The digestion mixture contained 1mg/mL collagenase II in DMEM with 0.1% BSA, 50 Units/mL DNase I and 2mM $CaCl_2$. WAT and digestion mixture was placed in an orbital shaker at 37˚C for 60 minutes. Digested tissue was then passed through 70μM cell strainers and centrifuged at 400xg for 5 minutes. Centrifugation resulted in the separation of the SVF cells from floating adipocytes. Adipocytes were removed, and SVF was resuspended in MACS buffer (PBS containing 0.5% BSA and 2 mM EDTA). Macrophages were isolated from the SVF by magnetic cell isolation technology using CD11b microbeads (Miltenyi Biotec), according to the manufacturer's protocol. Isolated cells were centrifuged and resuspended in DMEM supplemented with 10% FBS, 1 U/mL penicillin, 0.1 mg/mL streptomycin and 25 μg/mL Amphotericin B and 10% M-CSF derived from L929 cells before plating at the required cell densities. Cells were left to adhere overnight before assays were performed the following day. Flow cytometry confirmed ATMs as a macrophage population by analysing F4/80 and CD11b expression **(Supplementary Figure 9a).**

***Analysis of RNAseq dataset***

We utilised data from a previously described [8] HFD *in vivo* experiment in which adipose tissue macrophages were isolated from high-fat diet-fed mice and control diet-fed mice and RNA-seq data was generated. We analysed this dataset to examine the differential expression of M1 marker genes in adipose tissue macrophages as well as other genes of interest. We accessed this data from the NCBI Gene Expression Omnibus using GEO ID GSE158627. The $\log_2$ fold-change and p-value significance data were downloaded and analysed using the statistical computing language R [9] and the R package ggplot2 [10] for visualisation.

**Supplementary Table 1: qPCR primer sequences for murine and human genes**

| Primer | | Sense | | | Antisense | |
|---|---|---|---|---|---|---|
| **MURINE GENES** | | | | | | |
| *RPS18* | 5’ | CTTAGAGGGACAAGTGGCG | 3’ | 5’ | ACGCTGAGCCAGTCAGTGTA | 3’ |
| *PROCR* | 5’ | CAGGACACTTGTGTGGAGTT | 3’ | 5’ | CCAGGACCAGTGATGTGTAAG | 3’ |
| *F10* | 5’ | GAAGAGCAGAACTCAGTGGTGTG | 3’ | 5’ | CAAGCTCACTGTCGCTGGTGTT | 3’ |
| *F3* | 5’ | GCACCGAGCAATGGAAGAGTTTC | 3’ | 5’ | CTTTCTGTCCCGCTCGGTTCTT | 3’ |
| *SERPINC1* | 5’ | TCCGACGCATTCCACAAA | 3’ | 5’ | GGCCAGTAATCACGACAGAA | 3’ |
| *F7* | 5’ | CAGCCCACTGCTTCGATAATA | 3’ | 5’ | GTGTCACCCGTCGTACTTG | 3’ |
| *F11* | 5’ | GATTACCAAGCACACGCATTAC | 3’ | 5’ | AAGGACGGATGGCAGAATAC | 3’ |
| *F9* | 5’ | GCATTCTGTGGAGGTGCCATCA | 3’ | 5’ | TCTCCTTTGTTCTGTGTCTTCCTT | 3’ |
| *THBD* | 5’ | GGAGAATGGTGGCTGTGAGTAC | 3’ | 5’ | GCACGATTGAACCACAGGTCTTG | 3’ |
| *TFPI* | 5’ | AGGGAACGAGAACCGATTTG | 3’ | 5’ | TGCCTTCACAGCTGTCTTC | 3’ |
| *PROS1* | 5’ | TGGCAAGGAGACAGGTGTCAGT | 3’ | 5’ | GAGCAGTGGTAACTTCCAGGAG | 3’ |
| *F2* | 5’ | ACCTTGGGACTGTGAATGTC | 3’ | 5’ | GATGGGTGGTGGAGTTGATT | 3’ |
| *F8* | 5’ | GGCGAGTAGAATGCCTTATTGGC | 3’ | 5’ | ATCACGGATGCTTCCAGAAGCC | 3’ |
| *F5* | 5’ | TGATGCTGTCCAGCCCAATAGC | 3’ | 5’ | CGATCAAGCCTGAGTGGATGTC | 3’ |
| *SERPINE1* | 5’ | CCTCTTCCACAAGTCTGATGGC | 3’ | 5’ | GCAGTTCCACAACGTCATACTCG | 3’ |
| *SERPINB2* | 5’ | ACCCAGAGAACTTCAGTGGCTG | 3’ | 5’ | GAGAGAGGAGAAGGCTGAATGG | 3’ |
| *SERPINF2* | 5’ | TGCTTCTCCTCAACGCCATCCA | 3’ | 5’ | ATGTCCACCGACACTGTGAACC | 3’ |
| *PLAUR* | 5’ | GAGCACAGAAAGGAGCTTGA | 3’ | 5’ | GAAAGGTCTGGTTGCTATGGA | 3’ |
| *CCL3* | 5’ | GGAAGATTCCACGCCAATTC | 3’ | 5’ | TCTGCCGGTTTCTCTTAGTC | 3’ |
| *PLGrkt* | 5’ | CCCGCTAAGCTTCATCTTCA | 3’ | 5’ | CAGCTCCAGCTTCGTCTTT | 3’ |
| *ANXA2* | 5’ | GCAGAGGATGGCTCAGTTATT | 3’ | 5’ | CGTCGGTTCCTTTCCTCTTC | 3’ |
| *S100A10* | 5’ | GGCGACAAAGACCACTTGA | 3’ | 5’ | CTGGTCCAGGTCCTTCATTATTT | 3’ |
| *LRP1* | 5’ | TCCCAATGACAAGTCGGATG | 3’ | 5’ | GGCCCATATCCACCCAATAA | 3’ |
| *TFGB1* | 5’ | TGATACGCCTGAGTGGCTGTCT | 3’ | 5’ | CACAAGAGCAGTGAGCGCTGAA | 3’ |
| *PLAU* | 5’ | CTGCTTCATTCAACTCCCAAAG | 3’ | 5’ | GGCTGTCTTCCCTGTAGTATTC | 3’ |
| *CD11b* | 5’ | CAGGAGGCAAAGGCTGTTA | 3’ | 5’ | GACAGGCCCAAGGACATATT | 3’ |
| *TEK* | 5’ | GAACTGAGGACGCTTCCACATTC | 3’ | 5’ | TCAGAAACGCCAACAGCACGGT | 3’ |
| *TNFa* | 5’ | GCCTCTTCTCATTCCTGCTT | 3’ | 5’ | TGGGAACTTCTCATCCCTTTG | 3’ |
| *IL1b* | 5’ | GGAAGCAGCCCTTCATCTTT | 3’ | 5’ | TGGCAACTGTTCCTGAACTC | 3’ |
| *YM1* | 5’ | TACTCACTTCCACAGGAGCAGG | 3’ | 5’ | CTCCAGTGTAGCCATCCTTAGG | 3’ |
| *Arg1* | 5’ | GGCAGAGGTCCAGAAGAATG | 3’ | 5’ | TCCACCCAAATGACACATAGG | 3’ |
| *EGR1* | 5’ | AGCGAACAACCCTATGAGCACC | 3’ | 5’ | ATGGGAGGCAACCGAGTCGTTT | 3’ |
| *Jun* | 5’ | ACTCGGACCTTCTCACGTC | 3’ | 5’ | CGGTGTAGTGGTGATGTGCC | 3’ |
| *JunB* | 5’ | TCACGACGACTCTTACGCAG | 3’ | 5’ | CCTTGAGACCCCGATAGGGA | 3’ |
| *JunD* | 5’ | GTTGGTTGCGTGTTGAGTGT | 3’ | 5’ | CCAAGGATTACGGAACAGGA | 3’ |
| *Fos* | 5’ | CTCTGGGAAGCCAAGGTC | 3’ | 5’ | CGAAGGGAACGGAATAAG | 3’ |
| *FosB* | 5’ | ACCAGCTACTCAACCCCAG | 3’ | 5’ | GGGTAAGTGTCTCTTCTCGGG | 3’ |
| *Fosl1* | 5’ | GAGACGCGAGCCGAACAAG | 3’ | 5’ | CTTCCAGCACCAGCTCAAGG | 3’ |
| *Fosl2* | 5’ | AGCCTCCCGAAGAGGACAG | 3’ | 5’ | AGGACATTGGGGTAGGTGAAG | 3’ |
| **HUMAN GENES** | | | | | | |
| *PROCR* | 5’ | GCTCAATGCCTACAACCGCACT | 3’ | 5’ | CGAAGTGTAGGAGCGGCTTGTT | 3’ |
| *THBD* | 5’ | AACGACCTCTGCGAGCACTTCT | 3’ | 5’ | CCAGTATGCAGTCATCCACGTC | 3’ |
| *F3* | 5’ | CAGAGTTCACACCTTACCTGGAG | 3’ | 5’ | GTTGTTCCTTCTGACTAAAGTCCG | 3’ |
| *RSP18* | 5’ | GCAGAATCCACGCCAGTACAAG | 3’ | 5’ | GCTTGTTGTCCAGACCATTGGC | 3’ |

### Supplementary Table 2: RNAseq analysis results for adipose tissue macrophages

| gene_id | WT | WT_HFD | log2FoldChange | pvalue | padj | gene_name |
|---|---|---|---|---|---|---|
| ENSMUSG00000053475 | 27.8470323 | 174.538755 | -2.64268194 | 2.31E-07 | 5.46E-05 | *Tnfaip6* |
| ENSMUSG00000022015 | 2.06523219 | 43.6327736 | -4.322273236 | 3.56E-07 | 7.61E-05 | *Tnfsf11* |
| ENSMUSG00000027398 | 4406.63882 | 18838.8215 | -2.095927875 | 9.36E-07 | 1.58E-04 | *Il1b* |
| ENSMUSG00000025383 | 2.06523219 | 32.9663581 | -3.919215005 | 1.06E-05 | 0.00109188 | *Il23a* |
| ENSMUSG00000027399 | 204.194981 | 770.888191 | -1.915948792 | 1.21E-05 | 0.00121315 | *Il1a* |
| ENSMUSG00000020826 | 4.12775141 | 37.814729 | -3.158256182 | 3.21E-05 | 0.00258634 | *Nos2* |
| ENSMUSG00000028965 | 12.3779203 | 64.9656021 | -2.380549137 | 9.69E-05 | 0.00585847 | *Tnfrsf9* |
| ENSMUSG00000037411 | 632.173924 | 1971.34445 | -1.6406006 | 1.15E-04 | 0.00671608 | *Serpine1* |
| ENSMUSG00000028602 | 11.346647 | 55.2688622 | -2.272056275 | 1.87E-04 | 0.00966887 | *Tnfrsf8* |
| ENSMUSG00000031444 | 779.646187 | 2250.61052 | -1.5292803 | 3.06E-04 | 0.01471492 | *F10* |
| ENSMUSG00000037613 | 22.6906605 | 82.4197334 | -1.855364649 | 5.97E-04 | 0.02373882 | *Tnfrsf23* |
| ENSMUSG00000075122 | 579.578922 | 1571.83881 | -1.439188796 | 6.96E-04 | 0.02682575 | *Cd80* |
| ENSMUSG00000024401 | 823.990993 | 2218.61129 | -1.42882519 | 7.15E-04 | 0.02732243 | *Tnf* |
| ENSMUSG00000026875 | 833.272464 | 2239.94411 | -1.426472862 | 7.26E-04 | 0.02769791 | *Traf1* |
| ENSMUSG00000000982 | 647.643043 | 1643.59468 | -1.343423222 | 0.00148735 | 0.04701164 | *Ccl3* |
| ENSMUSG00000018916 | 2.06523219 | 18.4212419 | -3.083973287 | 0.00159793 | 0.04895949 | *Csf2* |

### SUPPLEMENTARY FIGURE LEGENDS

**Supplementary Figure 1: Gene expression changes in treated BMDMs and no significant thrombin generation in FVII$^{def}$ plasma**. **(a)** BMDMs were polarised towards M1 (100 ng/mL LPS and 20 ng/mL IFN-$\gamma$) or M2 (20 ng/mL IL-4 and IL-13) for 6 hr. Established genetic markers of M1 and M2 phenotypes were checked by RT-qPCR**, (a)** *TNFa* and **(b)** *IL1B* for M1 polarisation and **(c)** *YM1* and **(d)** *Arginase1* for M2 polarisation. **(e)** BMDMs were polarised for 6 hr or treated with 5 µg/mL ssRNA for 12 hr before gene expression analysis. Heat map showing coagulation-associated gene expression profiles for M1, M2 and ssRNA-treated BMDMs. Data were normalised with respect to *RPS18* mRNA levels, and results expressed as fold change compared to untreated BMDMs. **(f)** CAT analysis was performed in presence of BMDMs polarised for 18 hr with either FVII-deficient plasma or FXII-deficient plasma and **(g)** lag-times, **(h)** peak thrombin and **(i)** ETP determined. A Student's t-test or One-Way ANOVA was used where appropriate to determine the statistical significance with *P≤0.05 and **P≤0.01 for 3 independent experiments measured in duplicate.

**Supplementary Figure 2: M1 polarised human macrophages have enhanced thrombin generation.** HMDMs were polarised towards M1 (100 ng/mL LPS and 20 ng/mL IFN-$\gamma$) or M2 (20 ng/mL IL-4 and IL-13) for 18 hr. M1 and M2 cells were characterised by measuring surface expression of CD14 for macrophages and **(a)** CD86 for M1 population and **(b)** CD206 for M2 population. *F3* mRNA levels were measured by RT-qPCR in **(c)** THP1-derived human macrophages and **(d)** primary HMDMs. **(e)** CAT analysis was performed in presence of HMDMs differentiated with human serum alone and **(f)** lagtime, **(g)** peak thrombin and **(h)** ETP determined. **(i)** CAT analysis was performed in presence of HMDMs differentiated with human serum and 50 ng/mL recombinant M-CSF and **(j)** lagtime, **(k)** peak thrombin and **(l)** ETP determined. A paired t-test or One-Way ANOVA was used where appropriate to determine the statistical significance with *P≤0.05, **P≤0.01 and ****P≤0.0001 for 4-6 independent experiments measured in duplicate.

**Supplementary Figure 3: TLR7/8 agonist ssRNA promotes procoagulant activity in myeloid cells. (a)** BMDMs were treated with ssRNA and CAT analysis was performed to determine **(b)** lagtimes. **(c)** Primary human monocytes were treated with ssRNA and CAT analysis performed to determine **(d)** lagtimes and **(e)** peak thrombin. A One-Way ANOVA was used to determine the statistical significance with *P≤0.05, **P≤0.01 and ***P≤0.001 for 3-6 independent experiments measured in duplicate.

**Supplementary Figure 4: TF-containing extracellular vesicle release from BMDMs is enhanced with ssRNA treatment, but not by M1 polarisation**. To measure TF activity in BMDM supernatants, CAT analysis was performed in the presence of 80 µL supernatants (in serum-free media). **(a)** BMDMs were polarised towards M1 or M2 phenotypes for **(a-b)** 1 hr, **(c-d)** 3 hours, **(e-f)** 6 hours and **(g-h)** 18 hrs after which time, supernatants were collected and CAT analysis was performed to measure lagtime and ETP. **(i)** BMDMs were treated with ssRNA for 24-48 hr after which time supernatants were collected and CAT analysis performed to measure **(j)** lagtime, **(k)** peak thrombin and **(l)** ETP. A One-Way ANOVA was used to determine statistical significance with $*P \leq 0.05$ for 3 independent experiments measured in duplicate.

**Supplementary Figure 5: Glycolysis inhibition reverses inflammation-mediated procoagulant activity in human and mouse myeloid cells. (a)** Following 3 hours of LPS exposure, M1 or M2 polarisation or 6 hours of ssRNA treatment, the XF Seahorse Glycolytic Rate Assay was performed on BMDMs to determine basal glycolysis defined as the glycolytic proton efflux rate (glycoPER). **(b)** Schematic diagram of where the glycolytic inhibitors (2-DG and TEPP-46) target the glycolysis pathway. **(c-d)** 2-DG pre-treated primary human monocytes were activated with LPS and CAT analysis performed to determine **(e)** lagtime. 2-DG pre-treated primary HMDMs were polarised and CAT analysis was performed to determine lagtime for **(f-i)** human serum differentiated HMDMs and **(j-m)** M-CSF differentiated HMDMs. **(n)** CAT analysis performed with 2-DG pre-treated ssRNA-treated BMDMs to determine **(o)** lagtime and **(p)** peak thrombin. **(q)** CAT analysis performed with 2-DG pre-treated ssRNA-treated human monocytes to determine **(r)** lagtime and **(s)** peak thrombin. A one-way ANOVA was used to determine the statistical significance with $*P \leq 0.05$, $**P \leq 0.01$ and $***P \leq 0.001$ for 3-6 independent experiments measured in duplicate.

**Supplementary Figure 6: Cell viability not affected by 2-DG treatment.** 2-DG-treated BMDMs were polarised towards an M1 phenotype. Cell viability was measured using LIVE/DEAD™ Cell Stain Kit. Results show a representative graph.

**Supplementary Figure 7: Successful siRNA knockdown of *SERPINE1* and *SERPINB2* in M1 and ssRNA-treated macrophages. (a)** BMDMs were transfected with *SERPINE1* siRNA or non-targeting (NT; both 20nM) siRNA before M1 polarisation. *SERPINE1* mRNA levels were determined using RT-qPCR**. (b)** Before ssRNA exposure, BMDMs were transfected with SERPINB2 siRNA or NT siRNA (both 20 nM). *SERPINB2* mRNA levels were determined using RT-qPCR. A Student's t-test was used to determine statistical significance with $**P \leq 0.01$ for 3 independent experiments measured in duplicate.

**Supplementary Figure 8: Enhanced EPCR expression in primary human macrophages.** HMDMs were polarised and mRNA levels for **(a)** *PROCR* and **(c)** *THBD* were determined using RT-qPCR after 6 and 18 hours. HMDMs were polarised before measuring surface expression of **(b)** EPCR and **(d)** TM by flow cytometry. Results are shown as the % of live cells expressing ECPR or TM. A Student's t-test or one-way ANOVA was used where appropriate to determine statistical significance with $*P \leq 0.05$, $**P \leq 0.01$, 2-4 independent experiments measured in duplicate.

**Supplementary Figure 9: Macrophage Characterisation and gating strategy for ATMs in HFD *in vivo* experiment.** Mice were fed either control diet (CD) or High-fat diet (HFD) for 12 weeks. Mice were sacrificed and white adipose tissue collected, and adipose tissue macrophages (ATMs) were isolated using CD11b magnetic microbeads. Flow cytometry analysis measured the surface expression of CD11b, F4/80, EPCR and TM. **(a)** ATMs were confirmed as macrophages by CD11b+ F4/80+ staining. Gating strategies for **(b)** EPCR- and **(c)**TM-positive populations are shown.

**Supplementary Figure 10: Gating strategy for DSS colitis *in vivo* experiment.** Colitis was induced in mice by adding 2% (w/v) DSS to drinking water for 5 days with normal drinking water acting as a control. Mouse lamina propria leukocyte populations were isolated and isolated leukocytes were analysed by flow cytometry. Gating strategies for $EPCR^+$ DCs, granulocytes and monocytes are shown.

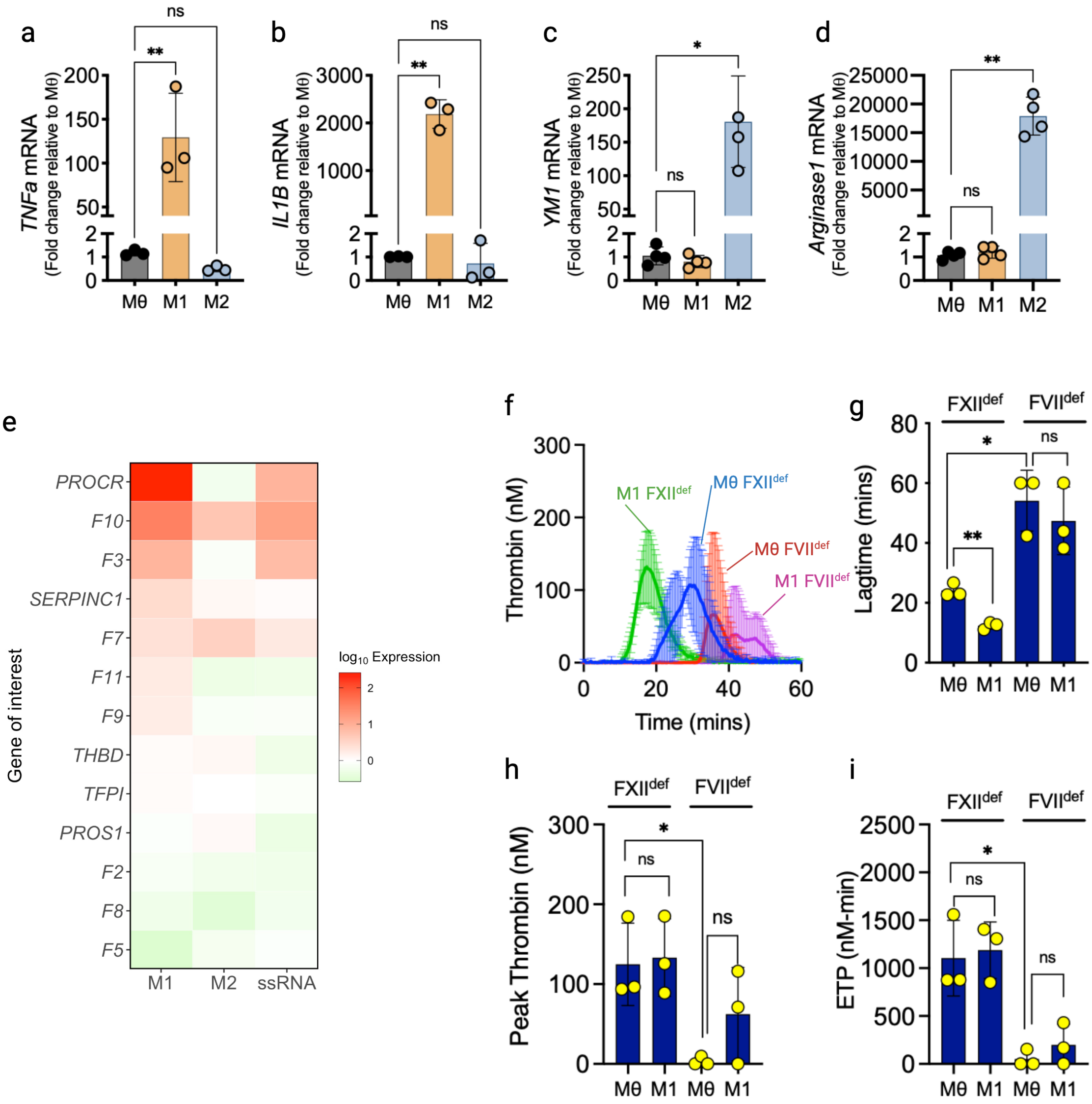

**Supplementary Figure 1**

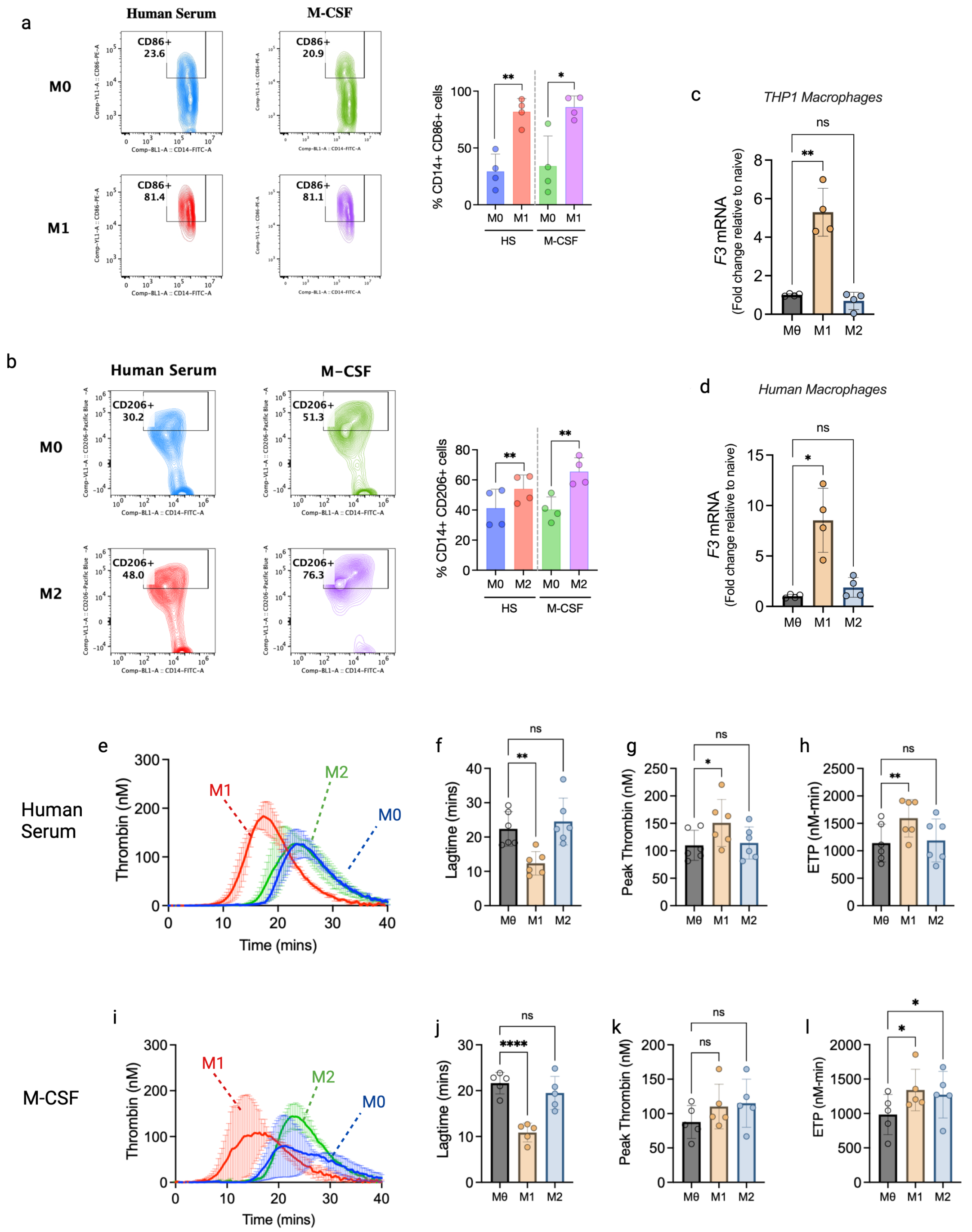

**Supplementary Figure 2**

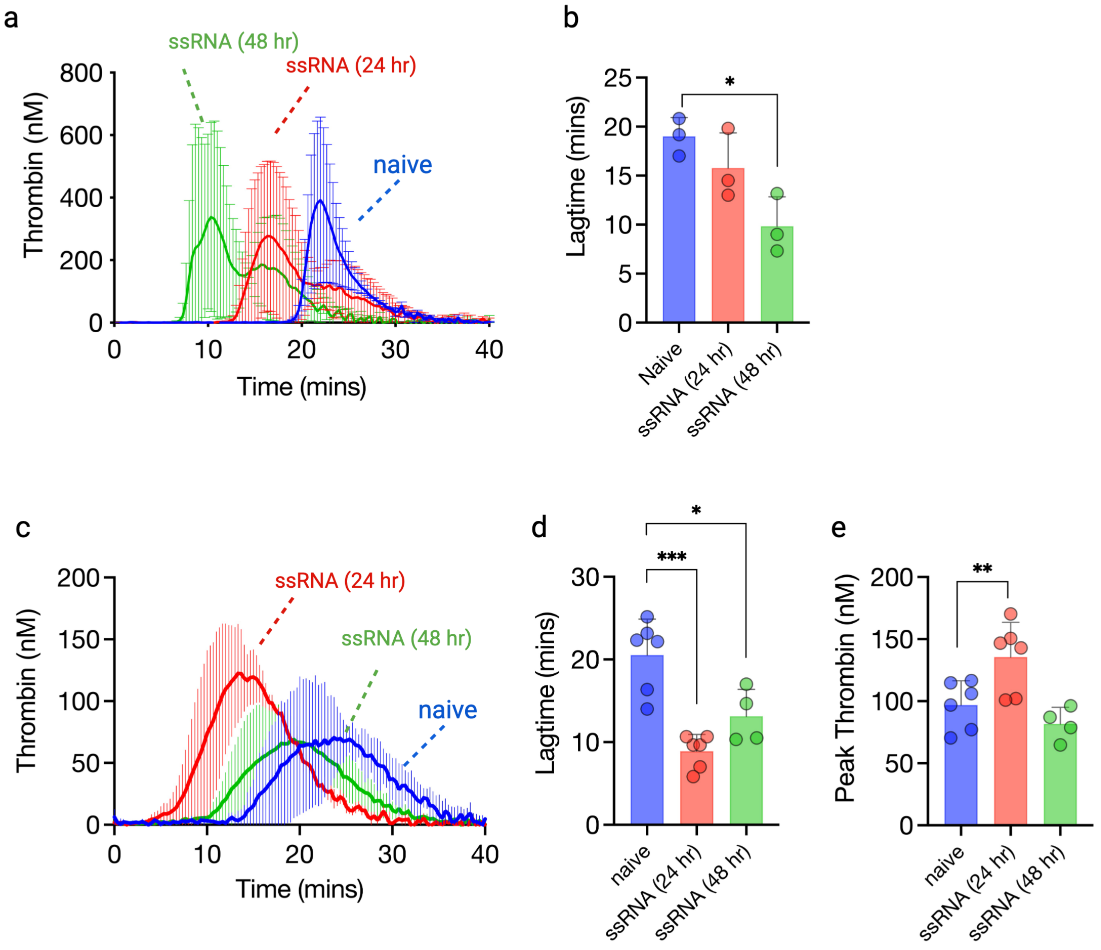

**Supplementary Figure 3**

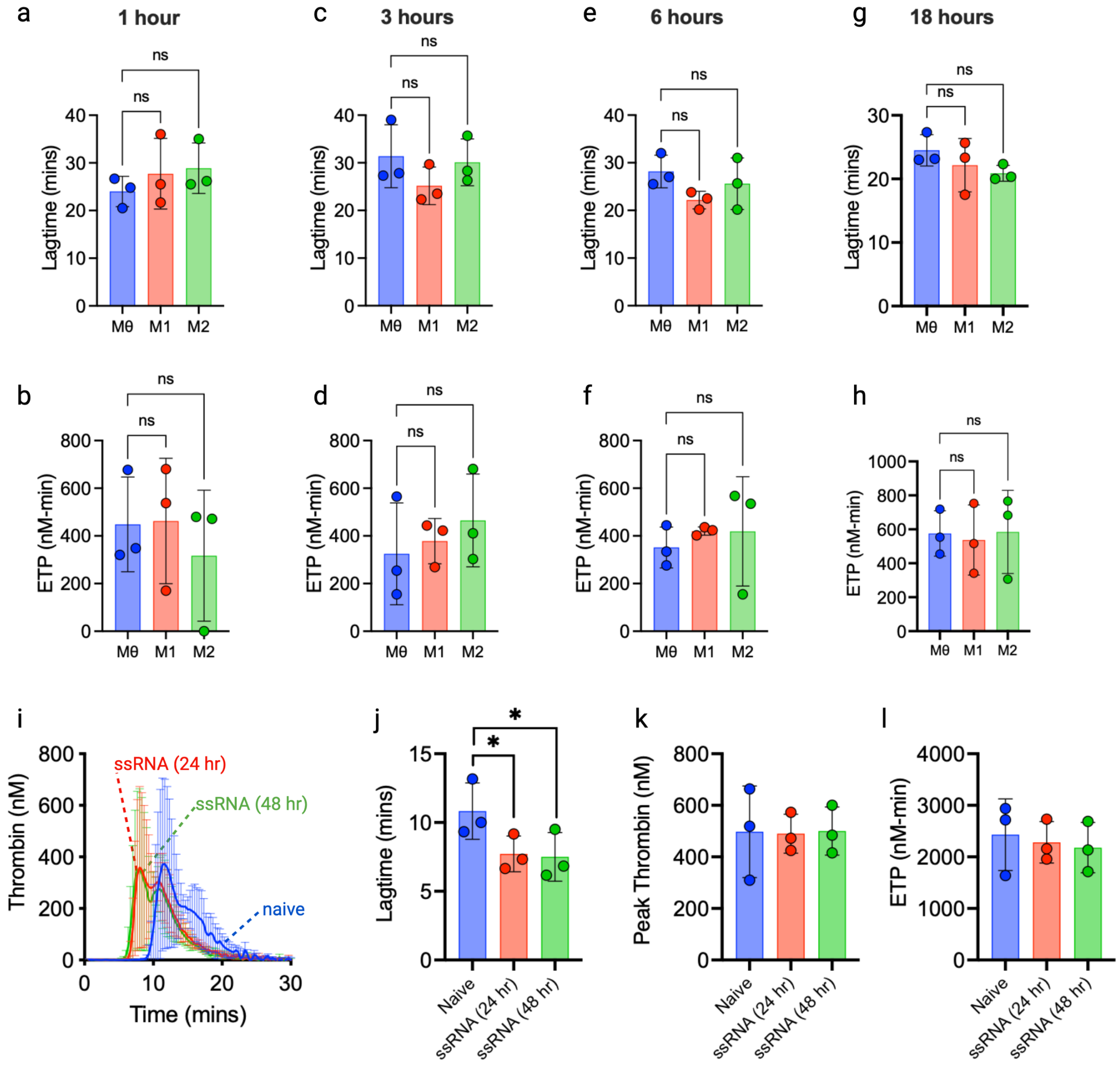

**Supplementary Figure 4**

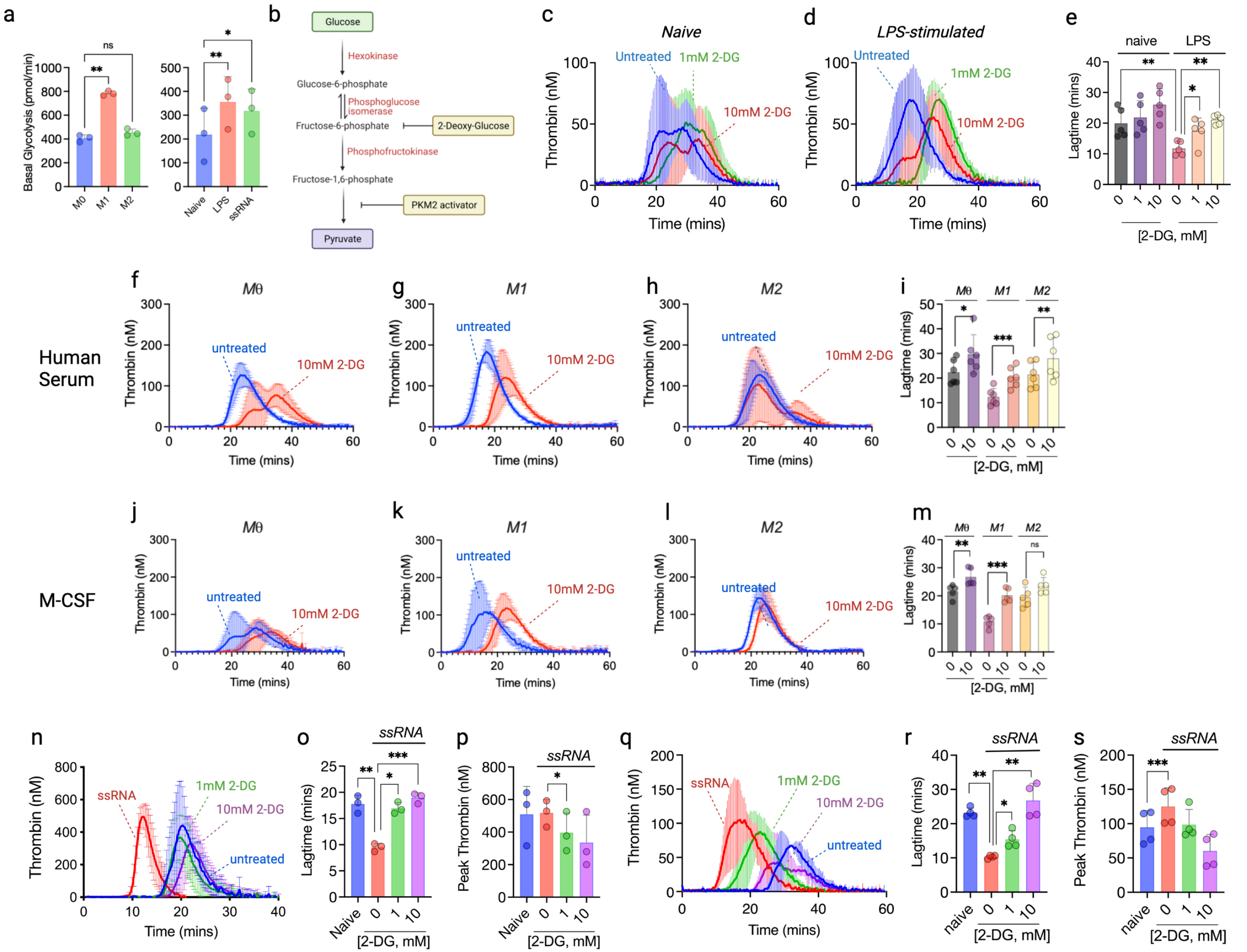

**Supplementary Figure 5**

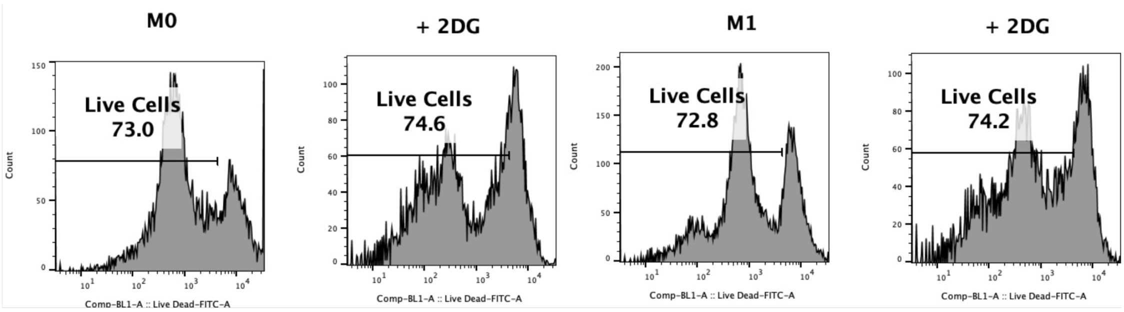

**Supplementary Figure 6**

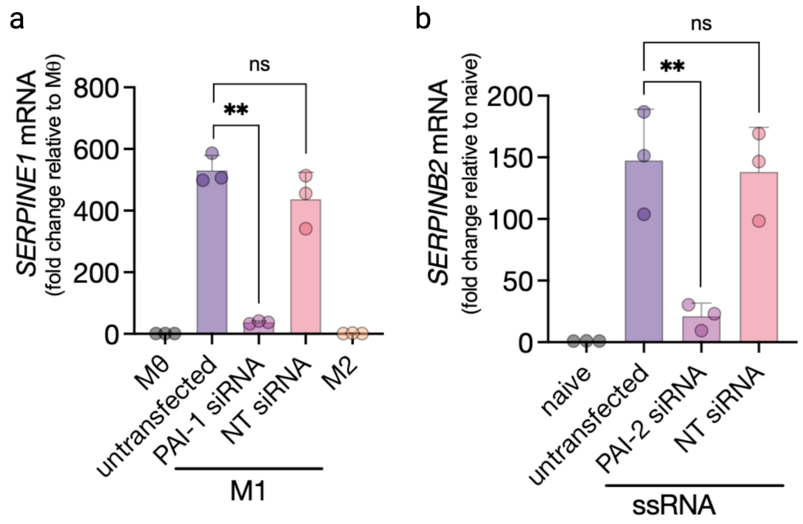

**Supplementary Figure 7**

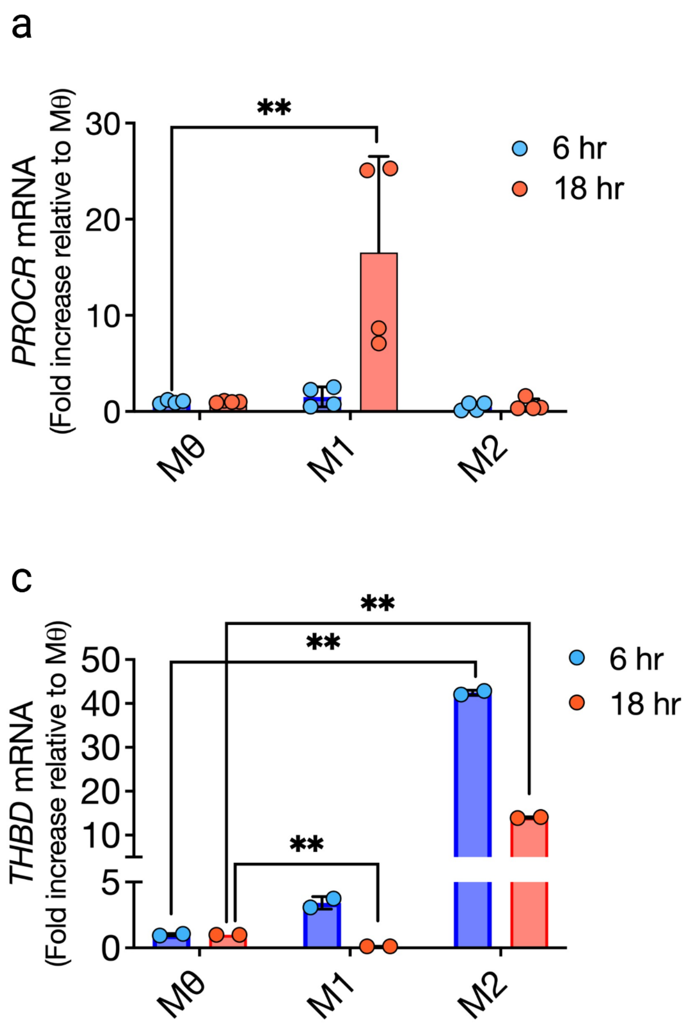

**Supplementary Figure 8**

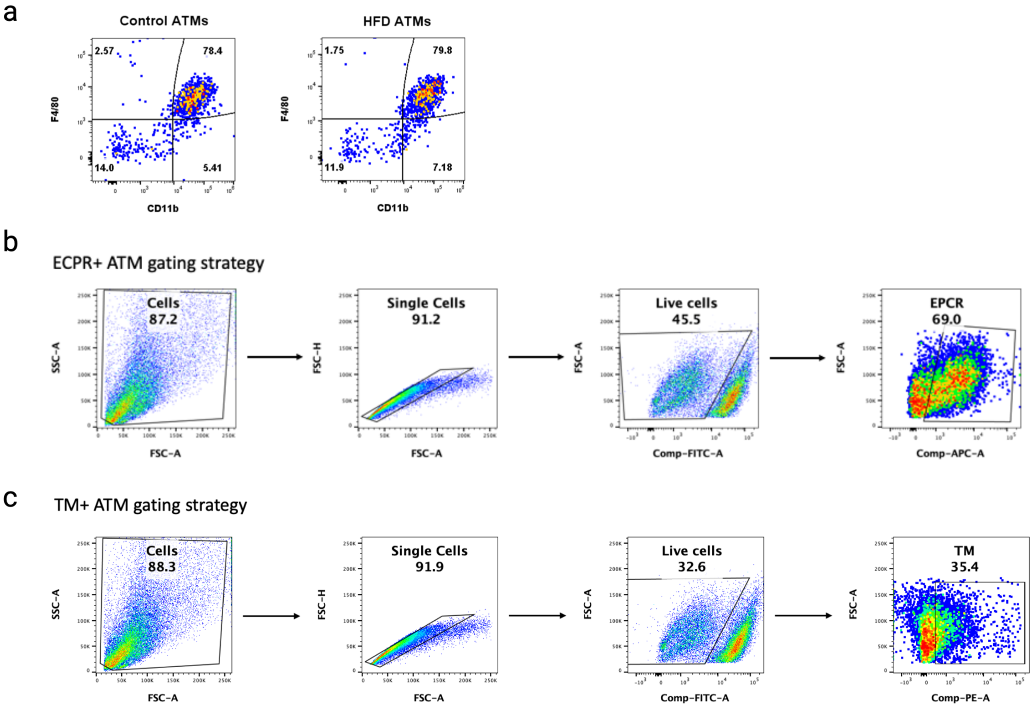

**Supplementary figure 9**

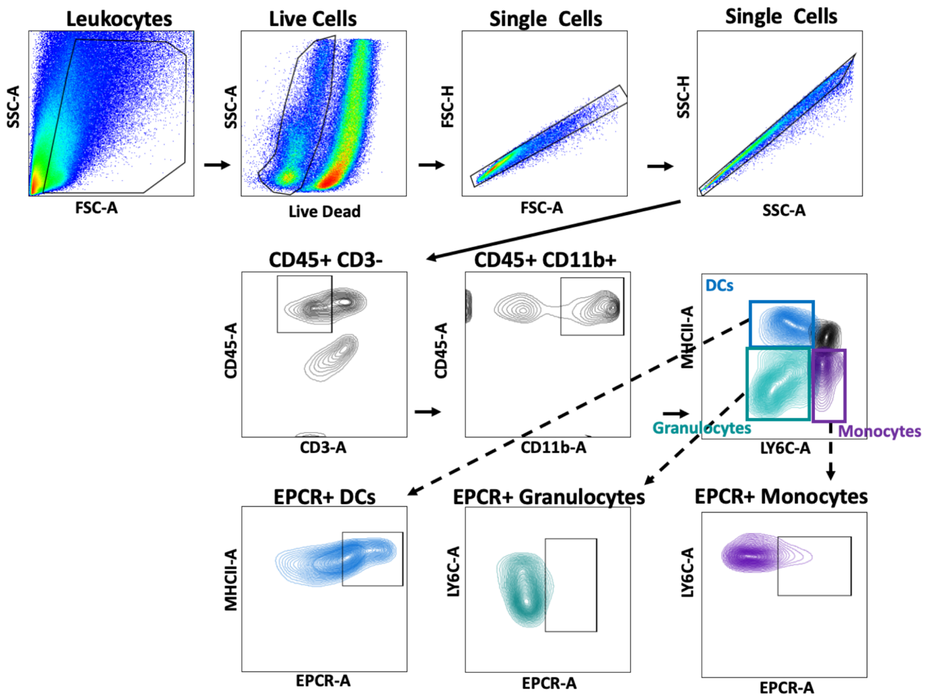

**Supplementary Figure 10**

9 R Core Team. R: A Language and Environment for Statistical Computing. Vienna, Austria: R Foundation for Statistical Computing; 2020.

10 Hadley, Wickham. ggplot2: Elegant Graphics for Data Analysis. Springer-Verlag New York; 2016.
